# Photoperiod and vernalization alleles and their combinations greatly affected phenological and agronomic traits in bread wheat under autumn and spring sowing conditions

**DOI:** 10.1101/2022.05.13.491906

**Authors:** Aduragbemi Amo, Dauren Serikbay, Luxing Song, Liang Chen, Yin-Gang Hu

## Abstract

The responses to temperature (vernalization) and day length (photoperiod) in wheat (Triticum aestivum L.) are entirely defined by allelic differences at the respective *VRN-1* and *PPD-1* loci. These two physiological factor greatly affect wheat phenology. Using molecular markers, a panel of 61 diverse bread wheat varieties were genotyped for *VRN-1* and *PPD-1* alleles, and the traits were evaluated under autumn and spring sowing conditions to investigate the impact of the alleles and their combinations on phenological stage, morphological traits, and yield traits. The results revealed that the earlier heading (HD), and flowering (FD) conferred by insensitive alleles for *VRN-1* and *PPD-1* genes was consistent regardless of the variation under the two sowing conditions. The effects of a single dominant allele of *VRN-1* were similar, with an additive effects when combined. The effects of a single *PPD-1* insensitive allele differed, with *Ppd-D1a* stronger than *Ppd-B1a,* and *Ppd-A1a* was the least, with an additive effects when combined. Secondly, varieties sown under spring condition recorded a larger flag leaf area when compared to those sown under autumn condition regardless of the single and combined influence at the *VRN-1* and *PPD-1* loci. When compared to other genes, *PPD-D1* had a consistent and significant effect on flag leaf area, spike length, and plant height across both sowing conditions. Furthermore, the sensitive allele *Ppd-D1b* had larger biomass across both sowing conditions, but this did not translate into increased yield because the *Ppd-D1a* insensitive allele recorded higher yield. The impact of *PPD-D1* gene on grain yield was evident across both sowing conditions. Overall, according to the findings of this study, *VRN-1* and *PPD-1* genes impact in modulating phenology stages, morphological, spike and yield traits was dependent on the sowing conditions and also on genetic background with the spring and insensitive alleles poised to be favored for selection.

## 1. Introduction

Bread wheat (Triticum aestivum L.) is extensively sown worldwide, with an average yield of over 700 million tons throughout the last decade (FAO, 2017) and it is an essential component of both energy and protein nutrition in human diet (Mahjourimajd et al., 2016). There will be 9.7 billion people on Earth by 2050, which means that agricultural productivity will have to expand by 25–70% (Hunter et al., 2017). This problem will only get more difficult in light of the projected effects of climate change. Therefore, more efforts should be put into enhancing the yield of wheat in areas where it is the primary source of calories and protein for humans. Every environment’s ability to increase plant yield potential can be maximized by optimizing water consumption; fertilizer utilization; radiation; and stress avoidance during the vegetative and grain-filling periods. Based on climate conditions, wheat production could be classified as autumn sowing, winter sowing and spring sowing, which requires specific types of wheat varieties to meet the growth and development and to achieve potential high yield. In China, for example, there are ten major wheat agroecological zones, and wheat varieties vary greatly in their adaptability to growing seasons, major biotic and abiotic stresses, and responses to temperature and photoperiod (Zhuang, 2003). Southern Australian wheat is sown in late autumn and early winter to avoid frost damage while still allowing enough time for the crop to reach maturity well before periods of drought (Pugsley 1983; Eagles et al., 2018). Wheat is cultivated in a roughly comparable manner in winter-rainfall regions of North Africa and West Asia (Hoshino and Tahir, 1987).

Vernalization (requirement for cold temperature) and photoperiod response are the most vital processes for wheat adaption to various environments. *VRN-1*, *VRN-2, VRN-3*, and *VRN-4* are the four series of vernalization genes, each with three copies across the three genomes (Distelfeld et al., 2009b; Kippes et al., 2014; Law et al., 1976; Pugsley 1972; Tan and Yan 2015 Yan et al., 2004b, 2006), while photoperiod genes (*PPD-1*) include *PPD-A1, PPD-B1,* and *PPD-D1* found on chromosomes 2 across the three genomes (Law et al., 1978). The regulation of phasic growth and acclimatization of wheat to specific environments is made possible due to *VRN-1* and *PPD-1* genes (Pugsley 1983; Trevaskis, 2010). These genes have been cloned, their functions have been clarified, and gene specific markers have been developed and used widely for wheat improvement (Nazim Ud Dowla et al., 2018). After the winter wheat vernalization requirement is met, photoperiod response will regulate the flowering timing, which is predominantly mediated by the *PPD-1* loci (Fjellheim et al., 2014). The photoperiod genes in wheat belong to the pseudo-response regulator family (Beales et al., 2007; Turner et al., 2005). The *VRN-1* and *PPD-1* allelic differences at the are mainly from mutations either in the coding region or the promotor regions of those loci, which resulted in the opposite effects, such as *PPD-D1b* for photoperiod-sensitive (long day) or *PPD-D1a* for photoperiod-insensitive (day-neutral) phenotype. It is possible to fine-tune phenological events prior to flowering by combining alleles of these genes without considerable modifications in anthesis date (González et al., 2011; Miralles and Richards, 2000; Slafer and Rawson, 1995; Slafer, 1996). It has been reported that combining *VRN-1* and *PPD-1* allelic variants have resulted in variations in phenological development such as flowering time and agronomic traits (Cane et al., 2013; Iqbal et al., 2006; Karsai et al., 2006) tillering, plant height, and spikelet number (Dyck et al., 2004; Miralles and Richard, 2000; Worland et al., 1998). Breeders use *VRN-1* and *PPD-1* allelic variants to modify crop phenology so that sensitive developmental phases take place in more optimal conditions.

The distribution of *VRN-1* and *PPD-1* allelic variants have been discovered to vary across continents, countries, and regions, with some haplotypes inadequately represented or absent in certain geographic areas (Kiss et al., 2014; Shcherban et al., 2014). This implies that different regions require different *VRN-1* and *PPD-1* combinations to meet local production conditions. In Afghanistan, the *PpD-D1a* characterized as vast majority (about 97%) of the wheat landraces studied (Manickavelu et al., 2014). The *VRN-A1, VRN-B1*, and *VRN-D1* dominant alleles were the most widespread in wheat cultivars across India’s various agro-climatic regions (Singh et al., 2013). Amongst the Japanese wheat cultivars, 83.8% possessed insensitivity for *Ppd-D1a* allele, but its frequency varied among genotypes from different geographical regions (Seki et al., 2011). Remarkably, the *Vrn-A1b* dominant allele is discovered to be widespread in wheat genotypes from Western Europe, North and South America (Efremova et al., 2016). Also, the dominant allele *Vrn-A1a* was widespread in spring cultivars cultivated in Northern Europe, western and central Europe but was least prevalent in southern Europe, while the *Vrn-B1a* allele is similarly represented in different regions in Europe and entirely dominates the *Vrn-B1c* allele primarily in Eastern European nations, such as Bulgaria, Romania (Shcherban et al., 2012).

Many studies have examined the effects of *VRN-1* and *PPD-1* allelic variants on the adaption and agronomic traits of wheat varieties or germplasm grown in various parts of the world (Gomez et al., 2014; Iqbal et al., 2007a; Santra et al., 2009; Shcherban et al., 2012; Yan et al., 2004a; Yang et al., 2009; Zhang et al., 2008), and many studies have examined the impact of these phenology genes on phenological traits and yield under different environments across different location constituting different longitude and latitude (Arjona et al., 2018, 2020; Chen et al., 2018; Eagles et al., 2010; Grogan et al., 2016; Shcherban et al., 2014; Royo et al., 2018). All these results confirmed that the rational utilization of *VRN-1* and *PPD-1* allelic variants in wheat breeding is key to the successful application of wheat varieties in their production regions. However, during wheat breeding, if one want to breed a variety for another region, sowing at different times (autumn or spring) in the same location can be a means to evaluate the adaptability of the lines in diverse environments. Therefore, understanding whether there are differences in the effects of the allelic variants of *VRN-1* and *PPD-1* genes on phenology and agronomical traits across autumn and spring sowing conditions, will be important for their proper utilization. Unfortunately, only few studies have been carried out in this regard (Steinfort et al., 2017; Ramírez et al., 2018).

Studying the effects of *VRN-1* and *PPD-1* allelic variants on plant phenology and agronomic traits will aid in determining which allele combinations are effective and beneficial in a specific growing area to assist plants in adapting their phenology to changing climate. Thus, promising cultivars with the right allelic combinations and observance of good agronomic practices are critical to increase productivity of wheat. Hence, the focus of this research is to analyze the influence of the allelic variants of *VRN-1* and *PPD-1* genes, as well as their combinations, on phenological development, main morphological traits, and yield traits under autumn and spring sowing conditions (SSC). In addition, we also explored the optimal combos of *VRN-1* and *PPD-1* alleles for specific growing environments, by using a wide panel of diverse wheat varieties, which were genotyped using the diagnostic markers of main *VRN-1* and *PPD-1* alleles, to promote proper selection coupled with the utilization of varieties in the right environments.

## 2. Materials and Method

### 2.1. Plant material and Experimental Field Setup

The diverse panel of 61 wheat varieties with different growth habits was formed, which mainly from various ecological regions of China, and some varieties from CIMMYT (Mexico), western USA, Australia, and northern and southern Kazakhstan were also included (Supplementary Table 1). These varieties were sown under Autumn and Spring conditions for two years at the experimental field of the Institute of Water Saving Agriculture in Arid Regions of China, Northwest A&F University, Yangling, Shaanxi, China (34°17′ N, 108°04′ E, elevation of 506 m).

The sowing date for the autumn sowing condition (ASC) for the two years was October 2 of 2019, October 8 of 2020, while that of the spring sown condition (SSC) was February 25 of 2020, January 30 of 2021, respectively. Field trials were conducted in randomized complete blocks with 3 replications, with each variety planted in three rows of 2.0 m length, 25 cm between rows, and 3.3 cm between plants. The experiment was irrigated especially for the spring sown conditions, and kept pest and disease free using normal agronomic practice.

### 2.2. Genotyping of *VRN-1* and *PPD-1* genes

A revised CTAB method was used to extract total genomic DNA from young leaves. (Priyadharshini et al., 2019). Allelic specific primers were used to detect the presence of dominant and recessive alleles of *VRN-A1, VRN-B1, VRN-D1, VRN-B3* genes as described by Steinfort et al. (2017), the primer set was show in (Supplementary Table 2) (Steinfort et al., 2017). The full length of *VRN-A1, VRN-B1, VRN-D1* genes were amplified for the recessive alleles and characterized as *vrn-A1, vrn-B1, vrn-D1*; while the ∼200 bp insertion of in the promoter of *VRN-A1,* and the deletion of the 1^st^ intron of both *VRN-B1* and *VRN-D1* genes, detected the dominant alleles, respectively, and were characterized as *Vrn-A1a*, *Vrn-B1a* and *Vrn-D1a* (Steinfort et al., 2017). The thermal cycling was performed as follow: 94°C for 2 min; 35 cycles of 94°C denaturation for 20s, 60°C - 62°C (dependent on the primer) annealing for 20 s, and 72°C extending for 30s; and 72°C for 10 min. Finally, PCR products were separated using 1.5-2% agarose gel electrophoresis. For *VRN-B3*, the primer set and PCR condition used was as described (Whittal et al., 2018).

For the *PPD-1* loci, the primer set and thermal cycling condition as described by (Seki et al. 2013) and (Beales et al., 2007, Díaz et al., 2012) was used to detect *PPD-A1* and *PPD-D1* genes respectively. For the detection of *PPD-B1* gene, SSR marker gwm148 was used as described (Nishida et al., 2013). These primer sets distinguishes the sensitive from the insensitive alleles. The sensitive allelic variants are characterized as: *Ppd-A1b*, *Ppd-B1b*, *Ppd-D1b*, and insensitive alleles as: *Ppd-A1a*, *Ppd-B1a*, *and Ppd-D1a*. The segments amplified and the product size of the *VRN-1* and *PPD-1* alleles investigated in this study was provided in **Table S2**.

### 2.3. Phenotypic Evaluation and derived variables

The crop growth stages were determined using Zadoks’ scale (Zadok et al., 1974). The days to heading (HD) and anthesis (FD) were estimated when half of the tillers had completed head emergence, anthers extruded from the florets and turned color, respectively.

The flag leaf length (FLL), Flag leaf width (FLW) after flowering and spike length (SL) were all measured using a metric ruler. The flag leaf area (FLA) was obtained by multiplying FLW x FLL x0.8 (correction for leaf shape) (Rebetzke and Richard, 1999). Plant height (PH), excluding awns, was evaluated briefly after flowering. Spikelets per spike (SPS) was determined by the average value of randomly collected sample spikes. Fifteen spikes were collected randomly, dried, weighted and threshed for measuring Grain number per spike (GNPS) and spike fertility (SF) was calculated as described in the methodology of Abbate et al. (2013). The grain number per spike (GNPS) was calculated by dividing the total of grains in the sample by the number of spikes. At maturity, a 0.5 m^2^ was also harvested for biomass (BIO), grain yield (GY), and thousand kernel weight (TKW) were measured and harvest index (HI) calculated.

### 2.4. Statistical analysis

The ANOVA, and mean comparisons of the *VRN-1* and *PPD-1* allelic variants and their combinations were performed using the “agricolae” package (LSD test at the 0.05 level) in the R software (R 3.4.1). Pearson’s coefficients of correlations of traits were computed using the “metan” package in R. The graphs and Box plots were created with ORIGIN software OriginPRO 2021 (OriginLab Corp., Northampton, MA).

## 3. Result

### 3.1. Characteristics of the autumn and spring sowing conditions

For autumn sowing conditions, the monthly average temperature ranged from 1.2°C to 23.7°C and 0.1°C to 25.2°C, and **t**he average temperature for the period of sowing to heading, and sowing to flowering were 6.8 ^◦^C and 6.2 ^◦^C, 7.7 ^◦^C and 8.2 °C for the 2019-2020 and 2020- 2021 growing seasons, respectively (Fig. 1), which could meet the vernalization requirement of most varieties, though the leaves of the spring varieties were frozen damaged over winter period. For spring sown conditions (SSC), the monthly average temperature ranged between 4.6°C to 23.7°C and 6°C to 25.2°C, and the average temperature for the period of sowing to heading, sowing to flowering, was 11.4 °C and 10.2 °C, 13.1 °C and 11.2 °C, for the 2020 and 2021, (Fig. 1), respectively, which also could meet the vernalization requirement of most varieties, while some strong winter varieties, especially those from southern Kazakhstan just had few tillers headed. As the temperature raised up quickly in the middle of June, the wheat plants of spring sowing were senescent in a few days, which could not be fully matured (Fig. 2). However, we found that lots of the winter varieties matured badly at the spring sowing in 2020, so the sowing date in 2021 was moved earlier in January 30^th^. Overall, the cropping cycle under ASC (8 months) was longer than those of SSC (5 months).

**Fig. 1.**
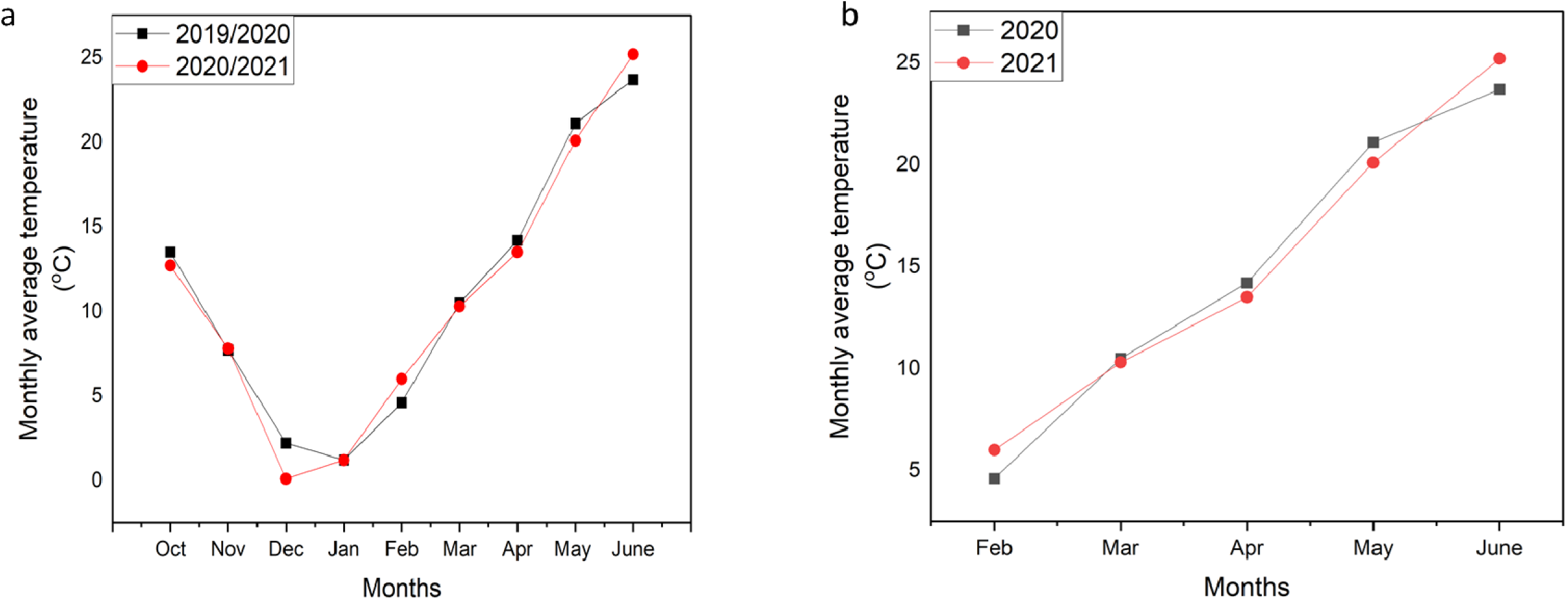
The distribution pattern of mean rainfall and temperature in each month under the two sowing conditions (a) autumn sowing conditions (b) spring sowing conditions

**Fig. 2.**
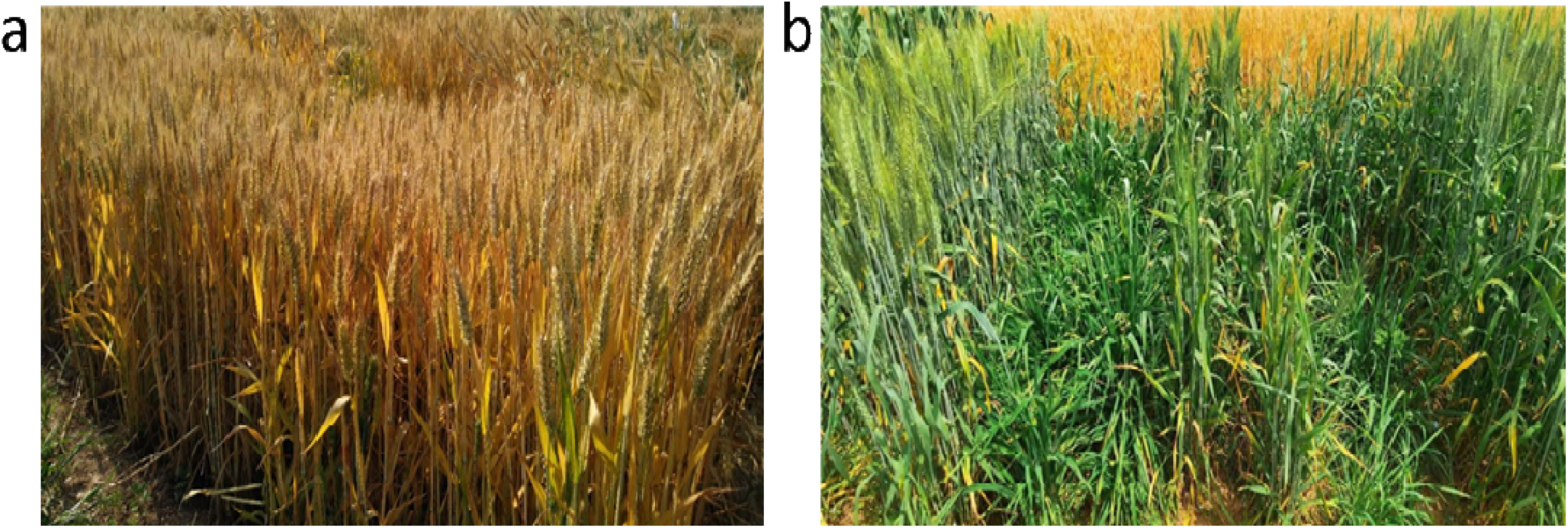
The variation in growth of varieties due to environmental changes under two sowing conditions (a) autumn sowing conditions (b) spring sowing conditions

### 3.2. Geographic Distribution of *VRN-1* and *PPD-1* alleles in the wheat panel

Genotyping with the markers of *VRN-1* and *PPD-1* revealed quite wide range of variations at the loci and their combinations among the 61 varieties. For the *VRN-1* genes, there was polymorphism observed for *VRN-A1, VRN-B1* and *VRN-D1* alleles (Fig. 3). The *VRN-D1* showed the largest variation with 38 varieties carrying the winter allele *vrn-D1* while 23 varieties carried the spring allele *Vrn-D1a* (Table 1). Followed by *VRN-A1*, with 44 varieties of winter allele *vrn-A1* while 17 varieties of spring allele *Vrn-A1a*. A low diversity was observed for *VRN-B1* gene as vast majority of the varieties carried the winter allele *vrn-B1* and only 7 varieties with the spring allele *Vrn-B1a*.

**Fig. 3.**
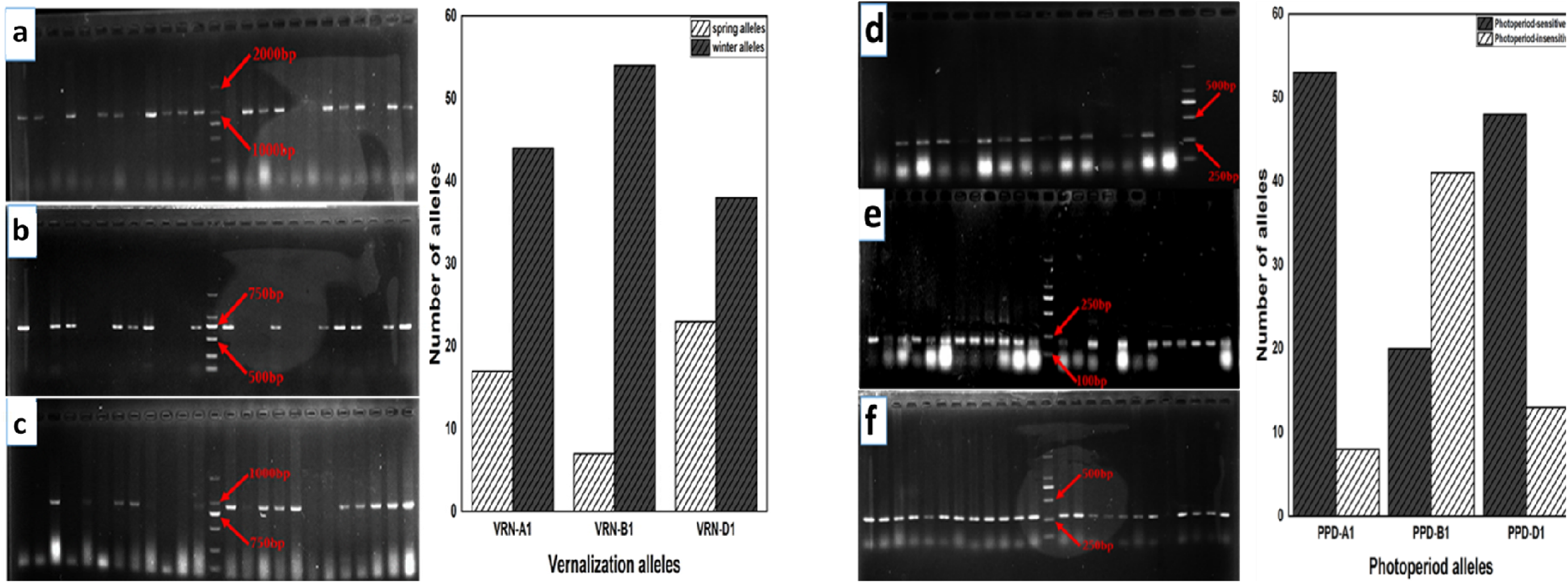
PCR amplification using genome specific primers for (a) *VRN-A1* (b) *VRN-B1* (c) *VRN-D1* (d) *PPD-A1* (e) *PPD-B1* (f) *PPD-D1* loci in different genotypes of wheat.

For *PPD-1* genes, *PPD-B1* showed the most allelic variation with 41 varieties carrying the insensitive allele *Ppd-B1a* while 20 varieties carried *Ppd-A1b*, the sensitive allele; followed by *PPD-D1* with 13 varieties carrying the sensitive allele *Ppd-D1b* while 48 varieties carrying the insensitive allele *Ppd-D1a*; the least was the *PPD-A1* as vast majority of the varieties (53) carried the sensitive allele *Ppd-A1b* while only 8 varieties carried the insensitive allele *Ppd-A1a*.

The geological distribution of the *Vrn-1* and *Ppd-1* alleles was quite consistent with their origin of those varieties. For the Chinese varieties, the winter alleles across the *VRN-1* loci were predominant at a frequency of 69.6% (*vrn-A1*), 93.5% (*vrn-B1*), and 56.5% (*vrn-D1*), respectively while for *PPD-1* loci, *Ppd-A1b* (87%), *Ppd-B1a* (73.9%), and *Ppd-D1a* (89.1%) alleles constituted the majority (Supplementary Table 1). For the 9 varieties that originated from Kazakhstan, the winter and sensitive alleles constitute the majority in the order: *vrn-A1* (8), *vrn-B1* (9), *vrn-D1* (9), *Ppd-A1b* (8), *Ppd-B1b* (6), and *Ppd-D1b* (7) (Table 2). For the “other regions” comprising of Australia, western USA and Mexico, two varieties originated from each country. For the Australian wheat, there was no spring allele *Vrn-A1a*, *Vrn-D1a* and insensitive alleles of *Ppd-A1a* and *Ppd-B1a* were lacking. Mexican wheat varieties possessed no winter alleles *vrn-B1* and *vrn-D1*, and also lacked sensitive alleles *Ppd-A1b*, *Ppd-B1b.* However, varieties from western USA possess equal number of spring and winter allele while lacking *Ppd-A1a, Ppd-B1b,* and *Ppd-D1b*.

### 3.3. The effects of *VRN-1* and *PPD-1* on phenological stage

#### 3.3.1. Autumn sowing conditions (ASC)

The wheat varieties carrying *Vrn-A1a*, *Vrn-B1a*, and *Vrn-D1a* alleles significantly accelerated HD by 4.6, 4.4, and 3.6 days, respectively and also advanced FD significantly by 2.8, 2.6, and 2.5 days than the winter alleles, respectively (Fig. 4). For the *PPD-1* loci, *Ppd-B1a* and *Ppd-D1a* alleles significantly advanced heading by 3.1 and 7.7 days, and FD by 2 and 6.1 days, compared with their sensitive alleles, while there was no significant difference for the two alleles of *Ppd-A1*.

**Fig. 4.**
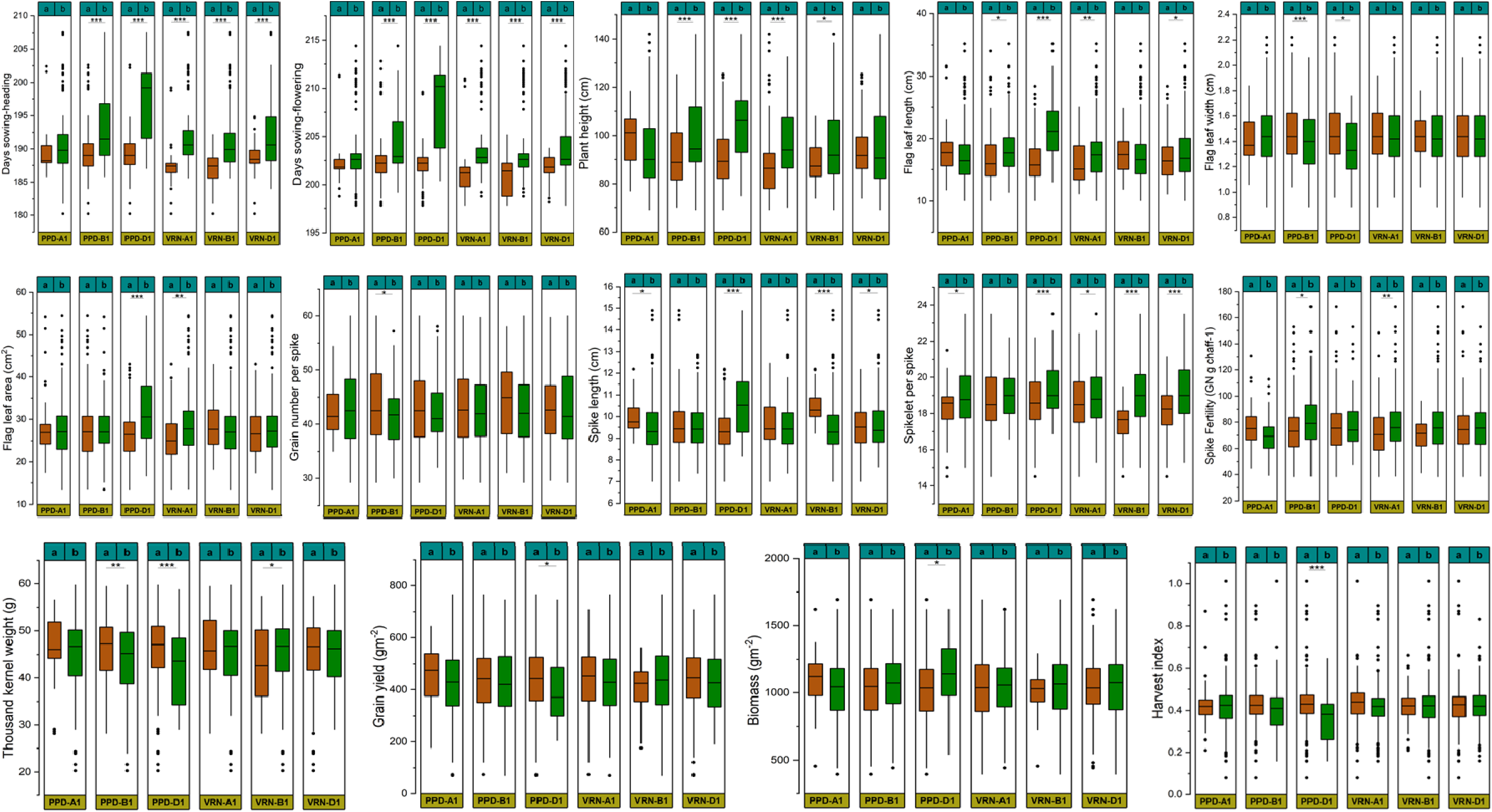
Box plots representing the frequency distribution of varieties in six genotypic groups (*VRN-A1, VRN-B1, VRN D1* and *PPD-A1, PPD-B1, PPD-D1*) for phenology stages, morphological, spike and yield traits observed in the field trials during autumn sowing conditions. a and b represent spring and winter alleles respectively at the *VRN-1* loci while representing insensitive and sensitive alleles respectively at the *PPD-1* loci respectively. *, **, and *** represent significant differences at 0.05, 0.01, and 0.001, respectively.

The two *VRN-1* allelic combinations revealed that the genotypes of double spring alleles of *Vrn-A1a+Vrn-B1a, Vrn-A1a+Vrn-D1a*, and *Vrn-B1a+Vrn-D1a* recorded earlier HD by 7.4, 6.7 and 6.4 days, and earlier FD by 4.4, 4.4 and 4.1 days than that of their double winter alleles, respectively (Table 3). The allelic combination of two *PPD-1* loci demonstrated that genotypes with the two insensitive alleles of *Ppd-A1a+Ppd-B1a, Ppd-A1a+Ppd-D1a,* and *Ppd-B1a+Ppd- D1a* favored early HD significantly by 5.5, 8.3 and 9.6 days, and earlier FD by 3.5, 9.6, and 7.3 days, than that of their sensitive alleles, respectively (Table 3). This suggested that there were additive effects for the *VRN-1* and *PPD-1* alleles.

According to the evaluation of *VRN-1* and *PPD-1* allelic combinations, genotypes with *Vrn-A1a+Ppd-A1a*, *Vrn-B1a+Ppd-B1a*, and *Vrn-D1a+Ppd-D1a* recorded earlier HD by 5.6, 6.8, and 9.1 days, and earlier FD 3.6, 4.3, and 6.8 days than that of the combinations of their wild types, respectively (Table 4). This suggest that there are interactive effects between the *VRN-1* and *PPD-1* alleles.

For three *VRN-1 or PPD-1* allelic combinations, the varieties with Vrn-A1a+Vrn-B1a+Vrn-D1a genotype recorded the earliest HD, and FD (184.3, and199 days respectively) and the varieties with *Ppd-A1a+Ppd-B1a+Ppd-D1a* genotype were associated with the earliest HD, and FD (Supplementary Table 3). Furthermore, the genotype *AdBdDdAsBiDi* recorded the earliest HD (182.5days) and FD (198.5days), and the genotype *ArBrDrAiBsDs* had the latest HD (203 days) and FD (211.5 days) (Supplementary Table 7).

#### 3.3.2. Spring sown conditions (SSC)

The wheat varieties carrying *Vrn-A1a*, *Vrn-B1a*, and *Vrn-D1a* alleles significantly accelerated HD by 4.7, 4.3, and 4.3 days, respectively and also advanced FD significantly by 2.9, 3.3, and 3.7 days than the winter alleles, respectively (Fig. 5). For the *PPD-1* loci, the photoperiod insensitive allele *Ppd-B1a*, and *Ppd-D1a* significantly advanced heading by 2.6 and 8.4 days, and FD by 2.5 and 8.0 days, compared with their sensitive alleles, while there was no significant difference for the two alleles of *PPD-A1*.

**Fig. 5.**
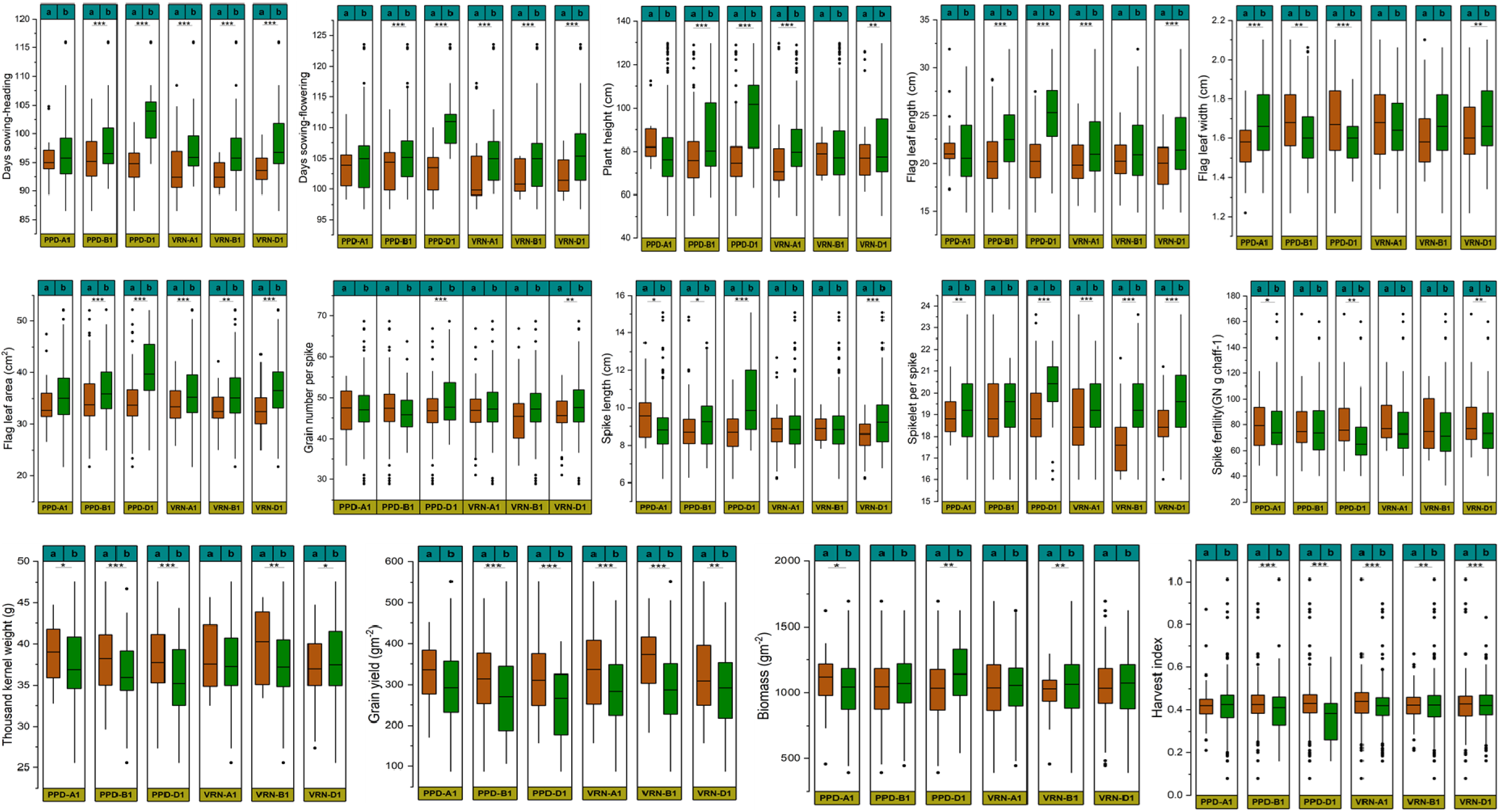
Box plots representing the frequency distribution of varieties in six genotypic groups (*VRN-A1 VRN-B1 VRN-D1*, and *PPD-A1, PPD-B1, PP-D1*) for phenology stages, morphological, spike and yield traits observed in the field trials during spring sowing conditions. a and b represent spring and winter alleles respectively at the *VRN-1* loci while representing insensitive and sensitive alleles respectively at the *PPD-1* loci respectively. *, **, and *** represent significant differences at 0.05, 0.01, and 0.001, respectively.

The two *VRN-1* allelic combinations revealed that the genotypes of double spring alleles of *Vrn-A1a+Vrn-B1a, Vrn-A1a+Vrn-D1a*, and *Vrn-B1a+Vrn-D1a* recorded earlier HD by 5.6, 6.4 and 7.4 days, and earlier FD by 4.6, 5.9 and 6.0 days than that of their double winter alleles, respectively (Table 3). The two *PPD-1* allelic combinations showed that genotypes with two insensitive alleles: *Ppd-A1a+Ppd-B1a, Ppd-A1a+Ppd-D1a* and *Ppd-B1a+Ppd-D1a* significantly promoted earlier HD by 5.2, 8.6 and 9.6 days, and earlier FD by 5.2, 8.2, and 9.2 days, than that of their sensitive alleles, respectively (Table 3). This suggested that there were additive effects for both the *VRN-1* and *PPD-1* alleles.

According to the evaluation of *VRN-1* and *PPD-1* allelic combinations, genotypes with *Vrn-A1a+Ppd-A1a Vrn-B1a+Ppd-B1a*, and *Vrn-D1a+Ppd-D1a* recorded earlier HD by 4.2, 6.2, and 9.9 days, and earlier FD 3.8, 5.2, and 9.2 days than that of the combinations of their wild types, respectively (Table 4). This suggested that there were interactive effects between the *VRN-1* and *PPD-1* alleles.

For the combined analysis of three *VRN-1 or PPD-1* alleles, the varieties with *Vrn-A1a*+*Vrn-B1a*+*Vrn-D1a* genotype recorded the earliest HD, and FD (95.3, and 106.6 days respectively) and the varieties with *Ppd-A1a+Ppd-B1a+Ppd-D1a* genotype were associated with the earliest HD, and FD (Supplementary Table 3). The genotype with the earliest HD, and FD respectively (89.4, and 98.5 days respectively) was revealed in AdBdDdAiBiDi (Supplementary Table 9).

### 3.4. The effects of *VRN-1* and *PPD-1* Alleles on morphological traits

#### 3.4.1. Autumn sowing conditions (ASC)

The plant height of wheat varieties with the *vrn-A1* and *vrn-B1* winter alleles was significantly taller by 10.9% and 7.0% than the spring ones, respectively whereas there was no significant difference for the two alleles of *VRN-D1*. The flag leaf length (FLL) was significantly shorter in varieties with spring allele *Vrn-A1a* and *Vrn-D1a by 11.0% and 7.7%* while there was no significant effect of *VRN-B1* recorded (Fig. 4). Furthermore, varieties with *Vrn-A1a* allele recorded a significantly reduced flag leaf area (FLA) by 11.5% while the alleles at *VRN-B1* and *VRN-D1* loci showed no significant effect (Table 4). Varieties with *Ppd-B1a* and *Ppd-D1a* alleles were significantly shorter in plant height (8.0% and 27.6% respectively) and FLL (7.0% and 30.3% respectively) when compared to the sensitive alleles. The FLA of insensitive allele *Ppd-D1a* was significantly smaller by 20.8% relative to the sensitive alleles. For flag leaf width (FLW), there was no significant difference observed for *VRN-1* alleles but *Ppd-B1a* and *Ppd-D1a* photoperiod sensitive lines significantly increased FLW by at least 3.6%.

The two *VRN-1* allelic combinations revealed that the genotypes of double winter alleles of *vrn-A1+vrn-B1, vrn-A1+vrn-D1, vrn-B1+vrn-D1* were significantly the tallest PH (97.5cm, 98.2, and 96.4cm respectively), longest FLL (18.6, 19.2, and 18.7cm respectively) (Table 5), and largest FLA (29.6cm^2^, 30.1cm^2^, and 29.2cm^2^ respectively) relative to their corresponding allelic combinations. However there was no significance difference observed FLW for the various *VRN-1* allelic combinations.

The two *PPD-1* loci allelic combination showed that the PH of genotypes with *Ppd-A1a+Ppd-B1b, Ppd-A1a+Ppd-D1b* and *Ppd-B1b+Ppd-D1b* were significantly the tallest (101.8cm, 112.5cm, and 107.4cm respectively) relative to their corresponding ones (Table 5). The varieties with *Ppd-A1a+Ppd-D1b* and *Ppd-B1a+Ppd-D1b* significantly had the longest FLL (26.1cm and 21.6cm respectively) and largest FLA (40.1cm^2^ and 34.5cm^2^ respectively) relative to their corresponding ones. Varieties with *Ppd-A1b+Ppd-B1a, Ppd-A1b+Ppd-D1a* and *Ppd-B1a+Ppd-D1a* genotypes significantly recorded the longest FLW (1.49cm, 1.50cm, and 1.49cm, respectively) when compared to their corresponding ones.

The evaluation of *VRN-1* and *PPD-1* allelic combinations shows that the PH of genotypes with *vrn-A1+Ppd-A1a, vrn-B1+Ppd-B1b* and *vrn-D1+Ppd-D1b* were the tallest (100.9cm, 100.1cm, and 106.1cm respectively), while *vrn-A1+Ppd-A1a* and *vrn-D1+Ppd-D1b* were associated with the longest FLL (18.3cm and 21.0cm respectively) relative to their corresponding ones (Table 6). Furthermore, *Vrn-A1a+Ppd-A1a* and *vrn-D1+Ppd-D1b* significantly had the longest FLA (29.9cm^2^ and 32.9cm^2^ respectively) when compared to the corresponding ones.

The combined analysis of three *VRN-1 or PPD-1* alleles indicated that varieties with *vrn-A1*+*vrn-B1*+*vrn-D1* was associated with the tallest PH (98.4cm) while *Vrn-A1a*+*Vrn-B1a*+*Vrn-D1a* recorded the longest FLL (20.9cm) and largest FLA (34.7cm^2^) (Supplementary Table 4). The varieties with *Ppd-A1a+Ppd-B1b+Ppd-D1b* genotype were associated with the tallest PH (112.5cm), longest FLL (26.1cm) and the largest FLA (40.1cm^2^). The AdBrDrAsBsDs genotype were the tallest PH (128.7cm) while ArBrDrAsBsDs recorded the largest FLA (40.1cm^2^). However, AdBdDrAsBiDi genotypes recorded the widest FLW (Supplementary Table 7).

#### 3.4.2. Spring sown conditions (SSC)

The varieties with *Vrn-A1a* and *Vrn-D1a* alleles were significantly shortened PH by 9.9% and 6.9% respectively and FLL by 6.9% and 9.5% respectively whereas *VRN-B1* gene showed no significant difference. (Fig. 5). The flag leaf area (FLA) was significantly shorter in varieties with spring allele *Vrn-A1a, Vrn-A1a* and *Vrn-D1a by 6.6%, 7.8%, and 12.6%*. Furthermore, varieties with *Vrn-D1a* allele recorded a significantly reduced flag leaf width (FLW) 3.7% while there were no significant differences observed for *VRN-A1* and *VRN-B1* genes (Table 4). Varieties with *Ppd-B1a* and *Ppd-D1a* alleles were significantly shorter in height (10.8% and 12.6% respectively) and FLL (9.3% and 18.5% respectively) when compared to the sensitive alleles. The FLA of insensitive alleles *Ppd-B1a* and *Ppd-D1a* was significantly smaller by 6.0% and 14.5% respectively, relative to the sensitive alleles. Furthermore, while *Ppd-B1b* and *Ppd-D1b* photoperiod sensitive alleles significantly reduced FLW by at least 3.7%, *Ppd-A1a* insensitive allele decreased FLW by 7%.

The two *VRN-1* allelic combinations indicated that genotypes of double winter alleles of *vrn-A1+vrn-B1, vrn-A1+vrn-D1, vrn-B1+vrn-D1* were significantly the tallest PH (86.3cm, 88.5, and 86.8cm respectively), longest FLL (22.5cm, 23.2cm, and 22.7cm respectively), and largest FLA (37.5cm^2^, 39.0cm^2^, and 38.2cm^2^ respectively) relative to their corresponding ones. Varieties with *Vrn-B1a+Vrn-D1a* genotype recorded the smallest FLW (1.53cm) when compared to other ones (Table 5).

The two *PPD-1* loci allelic combination showed that the PH of genotypes with *Ppd-A1b+Ppd-B1b, Ppd-A1a+Ppd-D1b* and *Ppd-B1b+Ppd-D1b* were significantly the tallest (88.1cm, 111.1cm, and 101.8cm respectively) and the longest FLW (22.7cm, 28.1cm, and 25.3cm respectively) relative to their corresponding allelic combinations (Table 5). The varieties with *Ppd-A1b+Ppd-B1b, Ppd-A1a+Ppd-D1b* and *Ppd-B1a+Ppd-D1b* significantly had the largest FLA (37.0cm^2^, 43.7 cm^2^ and 41.6cm^2^ respectively) relative to their corresponding allelic combinations. Varieties with *Ppd-A1b+Ppd-B1a, Ppd-A1b+Ppd-D1a* and *Ppd-B1a+Ppd-D1b* genotypes significantly recorded the longest FLW (1.71cm, 1.71cm, and 1.71cm respectively) when compared to their corresponding ones.

The evaluation of *VRN-1* and *PPD-1* allelic combinations shows that the genotypes *vrn-A1+Ppd-A1a, vrn-B1+Ppd-B1b* and *vrn-D1+Ppd-D1b* were significantly the tallest PH (86.9cm, 88.4cm, and 99.9cm respectively) while *Vrn-B1a+Ppd-B1b* and *vrn-D1+Ppd-D1b* were associated with the longest FLL (24.4cm and 25.5cm respectively) relative to their corresponding ones (Table 6). Furthermore, varieties with *vrn-B1+Ppd-B1b* and *vrn-D1+Ppd-D1b* genotypes significantly had the longest FLA (37.1cm^2^ and 41.1cm^2^ respectively) when compared to the corresponding ones.

The combined analysis of three *VRN-1 or PPD-1* alleles indicates that varieties with *vrn-A1*+*vrn-B1*+*vrn-D1* was associated with the tallest PH (86.6cm) and largest FLA (38.0 cm^2^) while *vrn-A1*+*Vrn-B1a*+*vrn-D1* recorded the longest FLL (24.4cm) (Supplementary Table 4). The varieties with *Ppd-A1a+Ppd-B1b+Ppd-D1b* genotype were associated with the tallest PH (111.1cm), longest FLL (28.1cm) and the largest FLA (43.7cm^2^). The tallest PH was the genotype AdBrDrAsBsDs while the PH of AdBrDrAsBsDi was the shortest. The largest FLA (43.7cm^2^) was observed in varieties with genotype ArBrDrAiBsDs (Supplementary Table 9).

### 3.5. The Effect of *VRN-1* and *PPD-1* Alleles on spike traits, yield and yield Components

#### 3.5.1. Autumn sowing conditions (ASC)

There was no significance difference observed for GY, biomass, GNPS, and HI amongst the different *VRN-1* alleles (Fig. 4). However, varieties with the *vrn-B1* winter allele increased TKW by 5.8% compared to the spring allele. The *Vrn-A1a* spring allele decreases SF by 10.6% while *vrn-B1* and *vrn-D1* winter alleles increase SPS. For *PPD-1* Loci, *PPD-D1* gene had a significant effect on GY, BIO, and HI with the *Ppd-D1a* insensitive allele significantly increasing GY and HI by 8.2% and 19.4% respectively. The effect of *PPD-B1* gene was significant for SF and GNPS with the *Ppd-B1a* allele increasing GNPS by 4.8% while *Ppd-B1b* increaseed SF by 7.6% (Table 6). In addition, varieties with *Ppd-D1b* sensitive allele recorded a 16.1% increase in SL.

The two *VRN-1* allelic combinations revealed that the genotypes of double spring alleles of *Vrn-B1a+Vrn-D1a* recorded the highest GNPS (47.8) when compared to other allelic combinations (Table 5). Interestingly, for GY, there was no significant difference amongst the different allelic combinations (Table 6). Varieties with the double spring alleles of *Vrn-A1a+Vrn-B1a, Vrn-A1a+Vrn-D1a, Vrn-B1a+Vrn-D1a* recorded the lowest number of SPS (17.1, 17.7, and 17.3 respectively) while *Vrn-A1a+Vrn-B1a and Vrn-B1a+Vrn-D1a* (10.7cm and 10.6cm respectively) relative to their corresponding ones. Varieties with *vrn-A1+vrn-B1, vrn-A1+Vrn-D1a* recorded the highest SF (77.4 and 79.7 respectively) whereas *Vrn-A1a+vrn-B1* and *vrn-B1+Vrn-D1a* had the highest TKW (46.3g and 46.4g respectively) in comparison to their corresponding ones.

The two *PPD-1* loci allelic combination showed that the genotypes of *Ppd-A1a+Ppd-D1b* and *Ppd-B1a+Ppd-D1a* recorded the highest GY (580.6gm^-2^ and 441.9gm^-2^ respectively) and HI (0.44 and 0.44 respectively) (Table 8). However, varieties with *Ppd-A1b+Ppd-B1b, Ppd-A1b+Ppd-D1b, and Ppd-B1b+Ppd-D1b* recorded the lowest TKW (43.2g, 41.8g, and 40.6g respectively) whereas varieties with genotypes of *Ppd-A1a+Ppd-D1b, and Ppd-B1b+Ppd-D1b* were associated with higher biomass (1329.8gm^-2^ and 1193.9gm^-2^ respectively) when compared to their corresponding ones. Furthermore, varieties with *Ppd-A1a+Ppd-D1a, and Ppd-B1a+Ppd-D1b* recorded the longest SL (11.0cm and 10.9cm respectively) and SPS (19.7 and 19.5 respectively) whereas varieties with *Ppd-A1b+Ppd-B1b, Ppd-A1b+Ppd-D1a, and Ppd-B1b+Ppd-D1a* recorded higher SF (80.0, 75.3, and 82.4) relative to their corresponding ones (Table 7).

The analysis of *VRN-1* with *PPD-1* allelic combinations showed that the genotypes of *Vrn-B1a+Ppd-B1b* and *vrn-D1+Ppd-D1b* recorded lower TKW (36.3g and 41.7g respectively) and HI (0.30, and 0.36 respectively) while varieties with *vrn-D1+Ppd-D1b* were associated with larger BIO (1150.1gm^-2^) relative to their ones (Table 9). However, there was no significant difference for the various allelic combinations on GY. Furthermore, varieties with *Vrn-A1a+Ppd-A1a, Vrn-B1a+Ppd-B1a* and *vrn-D1+Ppd-D1b* genotypes significantly had longer SL (10.6cm, 10.5cm, and 10.9cm respectively) when compared to the corresponding ones.

The three *VRN-1* loci combination analysis revealed that the various allelic *VRN-1* combinations had no significant effect on GY and SF (Supplementary Table 6). However, varieties with the genotype *Vrn-A1a*+*Vrn-B1a*+*vrn-D1* had the largest TKW (50.8g) and HI (0.48) whereas varieties with the genotype *Vrn-A1a*+*Vrn-B1a*+*vrn-D1* recorded the largest BIO (1164.4gm^-2^) relative to their corresponding ones. In addition, varieties with the genotype *Vrn-A1a*+*Vrn-B1a*+*Vrn-D1a* recorded the longest SL (11.0cm) while varieties with the genotype *Vrn-A1a*+*vrn-B1*+*vrn-D1* had the highest number of SPS (19.2) relative to the corresponding ones (Supplementary Table 5).

The three *PPD-1* loci combination analysis revealed that the various allelic combinations had no significant effect on GNPS (Supplementary Table 5). However, varieties with the *Ppd-A1a+Ppd-B1b+Ppd-D1b* genotype recorded the longest SL (11cm), highest SPS (19.7) but had the lowest SF (37.5). Furthermore, varieties with the *Ppd-A1a+Ppd-B1b+Ppd-D1b* genotype recorded the highest GY (580.6gm^-2^) and biomass (1329.8gm^-2^) (Supplementary Table 6). In addition, the genotype with the largest biomass (1522gm^-2^) and lowest HI (0.23) was AdBdDdAiBiDi (Supplementary Table 8). The GNPS was the highest with the AdBdDdAsBiDi (48) while the highest SF was recorded for ArBrDdAsBsDi.

#### 3.5.2. Spring sown conditions (SSC)

The single allelic effect of *VRN-1* loci revealed that the three *VRN-1* loci had a significant effect on GY, and HI with the spring alleles increasing GY and HI by at least 11.7% and 12.2% respectively (Fig. 5). However, there was no significant effect of these loci on biomass while only *VRN-B1* and *VRN-D1* had a significant effect on TKW. The effect of *VRN-A1* and *VRN-D1* genes on SPS and SF was significant with the spring alleles *Vrn-A1a* and *Vrn-D1a* increasing SF by 7.4% and 8.2% respectively. The significant effect of *VRN-D1* was observed for GNPS and SL with the winter allele *vrn-D1* increasing GNPS and SL (Fig. 5). For *PPD-1* loci, the effect of *PPD-B1* and *PPD-D1* were significant for GY and TKW with insensitive alleles increasing both traits in comparison to sensitive alleles. The effect of *PPD-A1* and *PPD-D1* genes on biomass was significant with *Ppd-A1a* and *Ppd-D1b* increasing biomass. Varieties with *Ppd-D1a* recorded a 37.5% increase in HI compared to *Ppd-D1b*. The significant effect of *PPD-1* was observed for GNPS, SL, SPS, and SF. The *Ppd-D1b* sensitive allele increases GNPS, SL, and SPS by 6.2%, 23.5%, and 6.9% respectively. However, *Ppd-D1b* resulted in a 10.6% decrease in SF.

The two *VRN-1* allelic combinations revealed that the genotypes of *Vrn-A1a+Vrn-B1a, Vrn-A1a+vrn-D1, and Vrn-B1a+vrn-D1* recorded significantly higher GNPS (47.9, 49.1, and 51.1 respectively) compared to their corresponding ones (Table 7). Varieties with the double winter alleles of *vrn-A1+vrn-B1, and vrn-B1+vrn-D1* recorded the longest SL (10.7cm, and 10.6cm respectively) and highest number of SPS (19.6, and 19.7 respectively) while genotypes of *Vrn-A1a+Vrn-B1a, Vrn-B1a+Vrn-D1a and Vrn-B1a+Vrn-D1a* recorded the highest GY (390.4gm^-2^, 336.3gm^-2^, and 369.3gm^-2^ respectively) relative to their corresponding ones (Table 8). Varieties with double winter alleles of *vrn-A1+vrn-D1, vrn-B1+vrn-D1* recorded the lowest SF (74.1 and 75.5 respectively) whereas *Vrn-A1a+Vrn-B1a, Vrn-A1a+vrn-D1* and *Vrn-B1a+vrn-D1* had the largest TKW (41.3g, 39.4g, and 40.5g respectively) in comparison to their corresponding ones.

The two *PPD-1* loci allelic combination showed that the genotypes *Ppd-A1a+Ppd-B1a, Ppd-A1a+Ppd-D1a and Ppd-B1a+Ppd-D1a* recorded the highest GY (378.1gm^-2^, 331.9gm^-2^, and 323.3gm^-2^ respectively) and HI (0.46, 0.45 and 0.45 respectively) (Table 8). However, varieties with *Ppd-A1a+Ppd-B1a, Ppd-A1a+Ppd-D1b, and Ppd-B1a+Ppd-D1a* recorded the largest TKW (40.27g, 42.7g, and 38.7g respectively) whereas varieties with genotypes of *Ppd-A1a+Ppd-D1b, and Ppd-B1a+Ppd-D1b* were associated with the highest biomass (990.3gm^-2^ and 825.3gm^-2^ respectively) compared to their corresponding ones. Furthermore, varieties with *Ppd-A1a+Ppd-B1b, Ppd-A1a+Ppd-D1a, and Ppd-B1a+Ppd-D1b* recorded the longest SL (10.9cm, 12.2cm and 10.9cm respectively) whereas varieties with *Ppd-A1a+Ppd-D1b, and Ppd-B1b+Ppd-D1a recorded the* highest SF (82.8, and 80.9) relative to their corresponding ones (Table 7).

The analysis of *VRN-1* with *PPD-1* allelic combinations showed that the genotypes *Vrn-B1a+Ppd-B1b* and *vrn-D1+Ppd-D1b* recorded the lowest TKW (35.2g and 35.9g respectively) while varieties with *Vrn-B1a+Ppd-B1b* and *vrn-D1+Ppd-D1b* were associated with the largest BIO (920.8gm^-2^ and 792.6gm^-2^) relative to their corresponding ones (Table 9). However, genotypes of *Vrn-A1a+Ppd-A1a, Vrn-B1a+Ppd-B1a* and *Vrn-D1a+Ppd-D1b* recorded the largest GY (336.2gm^-2^, 369.9gm^-2^, and 328.8gm^-2^). Furthermore, varieties with *vrn-A1+Ppd-A1a* and *vrn-D1+Ppd-D1b* genotypes significantly had the longest SL (9.8cm and 10.9cm respectively) compared to the corresponding ones.

The three *VRN-1* loci combination analysis revealed that varieties with *Vrn-A1a*+*Vrn-B1a*+*vrn-D1* had the highest GNPS (55), SF (93.8), and largest TKW (43.1g) (Supplementary Table 6). However, varieties with the genotype *vrn-A1*+*vrn-B1*+*vrn-D1* had the longest SL (9.6cm) and SPS (19.7) whereas varieties with the genotype *vrn-A1*+*Vrn-B1a*+*vrn-D1* recorded the largest BIO (920.8gm^-2^) relative to their corresponding ones. In addition, varieties with the genotype *Vrn-A1a*+*Vrn-B1a*+*Vrn-D1a* recorded the largest GY (409.7gm^-2^) and HI (0.51).

For three *PPD-1* loci allelic combinations, varieties with the *Ppd-A1a+Ppd-B1a+Ppd-D1a* recorded the highest GY (378.1 gm^-2^) and HI (0.46) (Supplementary Table 6) while varieties with the *Ppd-A1a+Ppd-B1b+Ppd-D1b* had the largest TKW (42.3g) and biomass (990.3gm^-2^). The varieties with the *Ppd-A1b+Ppd-B1a+Ppd-D1b* genotype recorded the highest GNPS (54.3) and SPS (20.8) (Supplementary Table 5) whereas varieties with the genotypes *Ppd-A1a+Ppd-B1b+Ppd-D1a* and *Ppd-A1a+Ppd-B1b+Ppd-D1b* gave the highest SF (87.4) and SL (12.2) respectively. Furthermore, the AdBrDdAsBsDi genotype recorded the highest SF (113.6), largest GY (447.6gm^-2^) and biomass (1003.9gm^-2^) whereas genotype AdBdDdAiBiDi had the lowest SPS (13.5) but gave the highest TKW (44 g) (Supplementary Table 10).

### 3.6. Correlation between phenology stages, morphological, yield and related traits

### 3.6.1. Autumn sowing conditions (ASC)

The number of days to flowering showed close association with days to heading (r=0.95) (Fig. 6a) while PH was associated with Days to flowering (r=0.44) and days to heading (0.45). GNPS was closely associated with SPS (r=0.31). Grain yield had a positive correlation with BIO (r=0.54), HI (r=0.67) and TKW (r=0.43) but recorded a negative correlation with PH (r=-0.19) and days to flowering (r=-0.09). HI showed positive correlation with TKW (r=0.37), but had no association with SF.

**Fig. 6.**
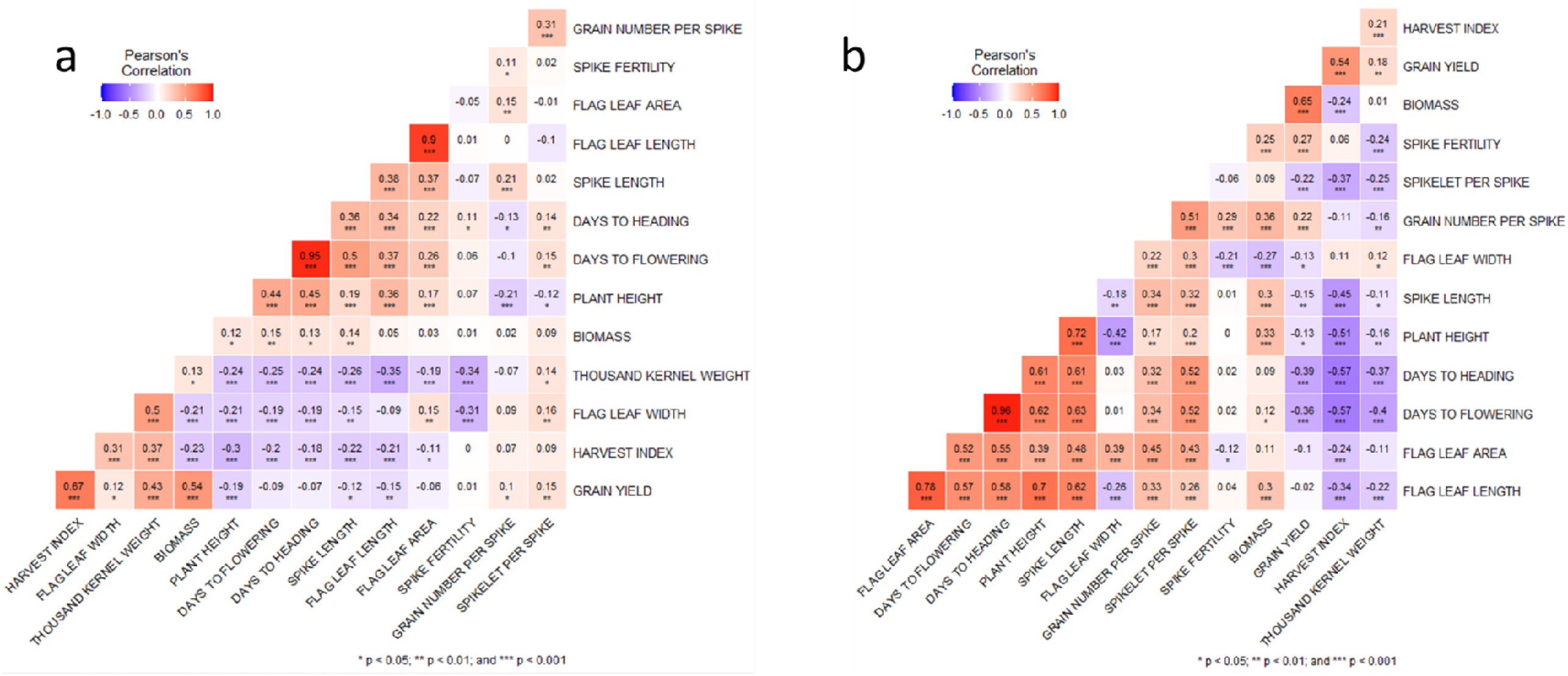
Pearson’s coefficient of correlations (above) and their significance probability (below) for pair-wise phenotypic traits observed in 2019-2020, 2020-2021 under (a) autumn sowing conditions (b) spring sowing conditions, respectively

#### 3.6.2. Spring sown conditions (SSC)

The Pearson’s correlation analysis indicated that days to flowering was positively correlated with days to heading (r=0.96), plant height (0.62), and spike length (0.63), but showed a negative correlation with HI, TKW and GY (Fig. 6b). GY was positively correlated with BIO, GNPS, SF while showing a negative correlation with the phenology stages days to heading (r=- 0.39), days to flowering (−0.36).

## 4. Discussion

### 4.1. Frequency distribution of *VRN-1* and *PPD-1* alleles

At the vernalization loci, more recessive (winter) *VRN-1* alleles were found using gene-specific molecular markers. This finding was in agreement with the study of Kiss et al. (2014) who observed and verified the predominance of the recessive alleles across the three *VRN-1* genes in wheat samples collected from Europe, Asia, America, Africa and Australia. Allelic variations in *VRN-1* genes were previously shown to have the strongest correlations with wheat genotypes’ geographic origins (Andeden et al., 2011; Yang et al., 2009; Zhang et al., 2008). It is estimated that the Yellow and Huai valley wheat production region (YHW) contributes 60–70% of the country’s wheat production despite its small size (Chen et al., 2018). There have been reports that the *VRN-A1* and *VRN-B3* loci are becoming scarcer and less common in the YHW region of China, which includes our experimental site (Chen et al., 2013; Zhang et al., 2008). In a similar study, *VRN-A1* and *VRN-B3* dominant alleles were found to be absent in wheat genotypes from the YHW in China (Chen et al., 2018). Although, this could be true of *VRN-B3* for this study as there was no polymorphism for this gene, nonetheless we found that the dominant spring allele *Vrn-A1a* accounted for 27.9% of the varieties studied, constituting by both local and exotic varieties **(Table S1)**. Admittedly, the dominant *Vrn-B3b* allele responsible for late flowering has been identified in landraces from China and Iran (Chen et al., 2013, Derakhshan et al., 2013). However, the recessive *Vrn-B3a* variant of T. durum is more common in the Ukrainian and Russian varieties (Muterko et al., 2016).

### 4.2. Duration of phenological Stages

Heading is an important phase in wheat development and *VRN-1* gene play important role in modulating this growth stage. In this present study, spring variants of *VRN-A1, VRN-B1*, and *VRN-D1* genes accelerated heading in both ASC and SSC. The effect of spring allele in promoting heading as revealed in this study is consistent with other reports. For example, it was discovered that the spring allele of the *VRN-D1* gene caused early HD in recombinant inbred lines, both in a controlled environment and in experiments conducted in autumn sowing fields (Kato et al., 2001) and Australian wheat genotypes (Eagles et al., 2010; Cane et al., 2013). Genotypes with more than one spring allele, such as those with two spring alleles (*Vrn-A1a+Vrn-B1a, Vrn-A1a+Vrn-D1a*, and *Vrn-B1a+Vrn-D1a*), headed significantly sooner than genotypes with single spring allele and this support the results of Iqbal *et al*. (2007*a*, 2007*b*) and Eagle et al. (2010).

Photoperiod genes play a profound impact on flowering in wheat and as expected varieties with photoperiod insensitive alleles heading and flowering earlier than the sensitive alleles across the two sowing conditions. The effect of photoperiod insensitive alleles in promoting flowering as reveled in this study is consistent with other reports. For instance in hexaploid wheat, on average, photoperiod insensitive genotypes reached anthesis earlier by 41.8 growing degree-days (Whittal et al., 2018) and 1.6 to 8 days earlier flowering (Grogan et al., 2016; Worland 1996) . The report of the significant effect of photoperiod insensitive alleles on flowering also hold true for durum wheat (Royo et al. 2015, Royo et al., 2016, Wilhelm et al., 2009). Interestingly several theories have tried to explain the phenomenon behind the early flowering conferred by insensitive alleles. Davidson et al. (1985) posited that the hastened anthesis was because of an acceleration in the development of the plant from emergence to floral initiation. Suppression of *PPD-1* and upregulation of *VRN-3* are linked to photoperiod insensitivity, and this promotes flowering via activating meristem identity genes (Corbesier et al., 2007; Yan et al., 2006). Another explanation to this is that it is also possible to induce earliness by reducing the amount of time it takes for heat units to accumulate between emergence and stem elongation (Foulkes et al., 2004). Genotypes with two insensitive alleles at the *PPD-1* loci (*Ppd-A1a+ Ppd-B1a, Ppd-A1a+ Ppd-D1a*, and *Ppd-B1a+ Ppd-D1a*) were found to head and flower more quickly at both sowing conditions than genotypes with only one sensitive allele or both sensitive alleles. Similarly Grogan et al. (2016) study reveals that both *PPD-B1* and *PPD-D1* insensitive alleles headed earlier than those with sensitive alleles at both loci.

Vernalization response genes are also known to contribute indirectly to yield by influencing flowering time (Goncharov, 2004; Stelmakh, 1993). The effect of the spring allele of *VRN-A1* in inducing early flowering in this study under ASC and SSC is consistent with other studies where the spring allele *Vrn-A1a* induces early flowering when compared to winter allele *vrn-A1* (Iqbal et al., 2007b; Kamran et al., 2013). From our study, the effect of photoperiod insensitive alleles on FD under ASC can be classified as *Ppd-D1a>Ppd-B1a>Ppd-A1a*, while under SSC, *Ppd-B1a* and *Ppd-A1a* have the similar effect but both were less than the effect conferred by *Ppd-D1a*, hence*: Ppd-D1a>Ppd-B1a/Ppd-A1a*. Our findings are in line with those of Worland et al. (1998), who found that the *PPD-D1* gene has the greatest effect, followed by the *PPD-B1* gene, with the *PPD-A1* gene having the least impact. The relative strength of these genes in bread wheat has been ranked differently in other studies. Photoperiod insensitivity alleles were classed by Shaw et al. (2012) as *Ppd-D1a > Ppd-A1a > Ppd-B1a* in terms of their effect on flowering date. According to Scarth & Law. (1984), *Ppd-B1a* may have an even stronger effect than *Ppd-A1a*, making it even more potent. Alleles conferring insensitivity in the B genome may have varying effects on earliness, according to Tanio and Kato. (2007), who discovered that the *Ppd-B1a* in hexaploid wheat had a greater effect than *Ppd-D1a*.

### 4.3. Morphological Traits

Varieties cultivated under ASC were taller than those under SSC and a plausible reason for this was that under SSC, full vernalization was not attained thus resulting in reduced stem elongation as opposed to wheat materials cultivated under ASC. In previous studies, the insensitive allele of *PPD-D1* gene has been shown to reduce plant height in wheat. For instance, accessions containing insensitive *Ppd*-*D1a* have plant height reduced by 15 % relative to sensitive *Ppd*-*D1b* accessions (Wilhelm et al., 2013), and similar effect was also observed in recombinant inbred lines (Gasperini et al., 2012). In this present study, Plant height was reduced by 13.9% in autumn sowing conditions when *Ppd-D1a* insensitive allele of were compared to sensitive allele. The impact of the *Ppd-D1a* allele was more pronounced in the SSC and accounted for about 27.7% reduction in plant height.

Admittedly, dwarfing genes are major genes that modulate plant height in wheat and have been utilized in wheat breeding programs culminating to the Green revolution (Hedden, 2003). However, the reducing effect of *Ppd-D1a* insensitive allele is analogous to dwarfing genes *Rht8*, *Rht15*, *Rht5* in wheat (Rebetzke et al., 2011; Tang et al., 2009; Wang et al., 2014; Zhao et al. 2021). Hence, since the effect of *Ppd-D1a* insensitive allele on promoting early heading in wheat is aimed at compensating the late heading of these dwarfing genes, therefore a combined selection for insensitive alleles of *PPD-D1* and these *Rht* genes was proposed (Chen et al., 2018; Zhang et al., 2019). Vernalization gene also had effect on plant height as seen from this present study but only alleles at *VRN-A1* loci showed consistent effect across the two sowing conditions. This finding is in agreement with the previous documentation of the role of *VRN-1* on plant morphology mainly relating to plant height, as seen in monosomic recombinant lines by Snape et al. (1985).

The flag leaf play a vital role in supplying assimilates for grain filling in wheat (Sanchez-Bragado et al., 2016). Individual leaf area have been reported to be responsive to the environment (Kirby et al., 1982; Dreccer et al., 2013) but not in the context of vernalization and photoperiod sensitivity. However, in this study we found that the FLA of varieties cultivated under SSC was higher than those in ASC. In addition, we found that *VRN-1* and *PPD-1* genes had significant effect on FLA, their sensitive alleles recorded higher leaf area than in insensitive alleles at both sowing conditions.

### 4.4. Spike and Yield Traits

In new wheat cultivars, *VRN-1* and *PPD-1* genes have a significant impact on targeting phase duration to improve yield potential (Fischer, 2011) and their effect on yield observed in this study showed varying effect at different sowing conditions. For instance *PPD-A1* gene had no effect on GY, TKW, HI at both sowing conditions but had an effect on BIO at SSC. The non-significant effect of *PPD-A1* gene on GY and TKW in this study is consistent with the findings of Arjona et al. (2018). However, the inverse was the case for other studies that recorded significant impact of *PPD-A1* on GY conferred by the insensitive alleles (Kamran et al., 2014; Maphosa et al., 2014; Slafer et al., 2005). A possible explanation for why alleles at *PPD-A1* had no impact on TKW and GY is that yield components are governed by multiple QTLs. (Wang et al., 2011), and by changing the length of the growth cycle, the *PPD-A1* gene acts as a modifier. The *PPD-B1* gene showed no effect on GY in ASC but impacted TKW at both sowing conditions with insensitive allele increasing TKW. This finding was in contrast to the studies that revealed that insensitive allele *Ppd-B1a* in both bread and durum wheat lowered TKW (Arjona et al., 2018; Maphosa et al., 2014). Studies have demonstrated that *PPD-D1* impact TKW, BIO, HI and GY (Worland et al., 1998; Foulkes et al., 2004; Guo et al., 2010), and also in this study we observed that *PPD-D1* had a significant impact on GY and TKW, BIO and HI at both sowing conditions.

The insensitive allele *Ppd-D1a* increased GY and TKW at both sowing conditions in comparison to the sensitive alleles, which is in support with findings that genotypes with the *Ppd-D1a* allele had a 13.5% higher yield than those with the *Ppd-D1b* allele (Whittal et al., 2018). *PPD-D1* allele yield differences have been reported in prior studies as well. For example, wheat genotypes bearing the photoperiod-insensitive *Ppd-D1a* gene have seen yields rise by up to 35% in southern Europe (Worland, 1996; Worland et al., 1998). Similarly in Germany, the *Ppd-D1a* allele improved yield by 7.7%, while in former Yugoslavia, the allele increased yield by 30% (Worland et al., 1998). The possible explanation of increased yield of *Ppd-D1a* genotypes lies in their ability to avoid the sweltering summer heat and dryness due to earlier flowering (Worland et al., 1998) and the same is observed in this study, and it also increases HI at both sowing condition. Sensitive alleles of *PPD-A1* and *PPD-B1* genes have shown to increase biomass (Royo et al., 2018) but in our study, the sensitive allele of *PPD-D1* gene recorded a significant rise in Biomass under both sowing conditions.

For spike morphology as relating to SL, *PPD-D1* showed consistence in its effect on SL at the two sowing conditions. This similar effect of *PPD-D1* gene on SL was also reported for wheat sown under vernalized condition (Steinfort et al., 2017) and this is synonymous to the ASC in this study as the wheat varieties underwent full vernalization. Overall, *PPD-D1* showed a consistent effect on FLA, SL, and PH across the two sowing condition compared to other phenology genes in this study.

Discussions on utilizing spike fertility (SF) as an indicator of wheat breeding success have been extensive (Abbate et al., 1998, 2013; Fischer, 2007, 2011; Foulkes et al., 2011; González et al., 2011; Lazaro & Abbate, 2012). It was discovered that this trait was moderately heritable with low GxE interaction and that was controlled by several genes (Martino et al., 2015; Mirabella et al., 2016). In comparison to the *Ppd-B1a* allele, the sensitive *Ppd-B1b* allele showed a 7.6% significant rise in SF under ASC. However, *PPD-D1* was found to have an impact on SF under SSC, with *Ppd-D1a* allele increasing SF by 10.6%. In a study examining the impact of photoperiod sensitivity on SF, the *PPD-B1* and *PPD-D1* genes’ insensitive alleles were found to be linked to an increase in SF (Ramírez et al., 2018).

Under ASC, *PPD-B1* played a significant role with insensitive allele contributing to 4.8% increase in GNPS in comparison to the sensitive allele. However, the effect of *PPD-B1* on GNPS was not consistence across two sowing conditions. It was reported that in certain environment, the effect of *PPD-B1* gene was found to be weaker when compared to *PPD-D1* (Worland et al., 1998). Interestingly, in some other studies, *Ppd-D1b* was observed to increase GNPS (González et al., 2005; Slafer et al., 2015), but *Ppd-D1a* also increased GNPS (Borner et al., 1993; Worland et al., 1998). However, in this study, *PPD-D1* had no effect on GNPS under ASC, but significantly impacted GNPS under SSC.

Regarding SPS, it had positive correlation with GY under ASC, but had a negative association with GY under SSC. However, although the *PPD-B1* has been shown to impact SPS in durum wheat (Arjona et al., 2018), there is no evidence that the *PPD-B1* gene has a direct effect on SPS in bread wheat., and in this study *PPD-B1* had no effect on SPS at both sowing conditions. On the other hand, *PPD-D1* gene significantly impacted SPS and was consistent across the two sowing conditions. The impact of *PPD-D1* gene on SPS as suggested may be due to its significant role in the determination of SPS by modulating *FLOWERING LOCUS T* (*FT*) expression at early growth phase, thereby concealing any lesser effects of *PPD-B1* gene (Arjona et al., 2018).

## 5. Conclusion

This study reveals the differential impacts of *VRN-1* and *PPD-1* alleles and their combinations at winter and spring sowing conditions, and while some alleles showed significant impact at one sowing condition, other alleles consistently impacted trait across the two sowing conditions. For the modulation of Phenology stages, insensitive allele at the *VRN-1* and *PPD-1* engender early HD, and FD independently of environmental and sowing conditions thus favoring the selection of such allele. For morphological and spike traits such as FLA, SL, and PH, *PPD-D1* gene had a significant impact compared to other phenology genes across both sowing season. Furthermore, only the insensitive allele *Ppd-D1a* significantly increased GY at the ASC but under SSC, both insensitive alleles *Ppd-D1a* and *Ppd-B1a* significantly increased GY.

## Acknowledgement

This work was financially supported by the Key Research and Development Program of Shaanxi Province (2021KWZ-23), the Construction of Overseas Agricultural Demonstration Park of NWAFU (HWJD202006), the China 111 Project (B12007), P. R. China.

