## supplemental table for "Photoperiod and vernalization alleles and their combinations greatly affected phenological and agronomic traits in bread wheat under autumn and spring sowing conditions"

**Table 1**

The distribution of vernalization and Photoperiodic genes in the bread wheat varieties surveyed.

| Genes | Alleles | N | Typical varieties | Description | References |
| --- | --- | --- | --- | --- | --- |
| VRN-A1 | vrn-A1 | 44 | Fengchan3, Beijing 411, Ningdong 10, Almaly | Full-length winter type, A genome. | Yan et al. 2004,  Steinfort et al. 2017 |
|  | Vrn-A1a | 17 | Xinong 585, Xiaoyan 22-3, Astana 2 | Promoter insertion, gene duplication. High activity: reduces vernalization requirement. |  |
| VRN-B1 | vrn-B1 | 54 | Dastan, Altyn masak, Zheng Mai 9023, Jimai 19 | Full-length winter type, B genome. | Fu et al. 2005,  Steinfort et al. 2017 |
|  | Vrn-B1a | 7 | Huai Mai 25, Xinmai 26, Clear white | Deletion 1st intron B genome. Common allele. Deletion: bp 1105-8885 |  |
| VRN-D1 | vrn-D1 | 38 | Jimai 22, Jimai 23, Drysdale, Quarrion | Full-length winter type, D genome | Fu et al. 2005,  Steinfort et al. 2017 |
|  | Vrn-D1a | 23 | Jimai 60, 87-341, Jimai 20, Changwu 134 | Deletion 1st intron D genome. Common allele |  |
| PPD-A1 | Ppd-A1a | 8 | Xinong 928, Zheng Mai 9023 | 1085 bp deletion in the promoter region | Wilhelm et al. 2009, Seki et al. 2013 |
|  | Ppd-A1b | 53 | Bulava, Steklovidnaya 24, Aifeng 3, Bainong 207, | Full length PPA1, A genome | Nishida et al. 2013, Seki et al. 2013 |
| PPD-B1 | Ppd-B1a | 41 | Fengchan 3, ning dong 11, xifeng 20, zhong mai 175 | 308 bp insertion in the promoter region | Nishida et al. 2013, Arjona et al., 2018 |
|  | Ppd-B1b | 20 | Xiaoyan 6, zhang 6878, xi nong 979, zhang han 58 | Full length PPB1, B genome |  |
| PPD-D1 | Ppd-D1a | 48 | Jimai 21, Jimai 22, Jimai 23, Jimai 44 | 2089 bp deletion in the promoter region | Diaz et al. 2012  Beales et al. 2007, Steinfort et al., 2017 |
|  | Ppd-D1b | 13 | Luohan 2, Ningchun 45 | Full length PPD1, D genome |  |

**Table 2**

The distribution of *VRN-1* and *PPD-1* alleles of wheat varieties used in this study based on country of origin

| Genes | Alleles | China | USA | Kazakhstan | Mexico | Australia |
| --- | --- | --- | --- | --- | --- | --- |
| VRN-A1 | vrn-A1 | 32 | 1 | 8 | 1 | 2 |
|  | Vrn-A1a | 14 | 1 | 1 | 1 | 0 |
| VRN-B1 | vrn-B1 | 43 | 1 | 9 | 0 | 1 |
|  | Vrn-B1a | 3 | 1 | 0 | 2 | 1 |
| VRN-D1 | vrn-D1 | 26 | 1 | 9 | 0 | 2 |
|  | Vrn-D1a | 20 | 1 | 0 | 2 | 0 |
| PPD-A1 | Ppd-A1a | 6 | 0 | 1 | 1 | 0 |
|  | Ppd-A1b | 40 | 2 | 8 | 1 | 2 |
| PPD-B1 | Ppd-B1a | 34 | 2 | 3 | 2 | 0 |
|  | Ppd-B1b | 12 | 0 | 6 | 0 | 2 |
| PPD-D1 | Ppd-D1a | 41 | 2 | 2 | 2 | 1 |
|  | Ppd-D1b | 5 | 0 | 7 | 0 | 1 |

**Table 3**

The effects of two allelic combinations of *VRN-1* or *PPD-1* loci on phenology stages in autumn sowing conditions (ASC) and spring sowing conditions (SSC)

| Allelic variations | N | Days to heading | | Days to flowering | |
| --- | --- | --- | --- | --- | --- |
|  |  | **ASC** | **SSC** | **ASC** | **SSC** |
| *vrn-A1+vrn-B1* | 41 | 192.5±0.3(a) | 97.9±0.3(a) | 204.2±0.2(a) | 105.5±0.3(a) |
| *vrn-A1+Vrn-B1a* | 3 | 189.7±0.4(ab) | 93.5±0.5(b) | 202.8±0.4(ab) | 102.6±0.5(b) |
| *Vrn-A1a+vrn-B1* | 13 | 188.5±0.4(b) | 94.4±0.6(b) | 201.9±0.3(b) | 102.9±0.6(b) |
| *Vrn-A1a+Vrn-B1a* | 4 | 185.1±0.4(c) | 92.3±0.5(b) | 199.7±0.3(c) | 100.9±0.5(b) |
| *vrn-A1+vrn-D1* | 29 | 193.6±0.4(a) | 99.0±0.4(a) | 205.0±0.3(a) | 106.5±0.4(a) |
| *vrn-A1+Vrn-D1a* | 15 | 189.8±0.2(b) | 94.9±0.2(b) | 202.4±0.1(b) | 103.1±0.3(b) |
| *Vrn-A1a+vrn-D1* | 9 | 188.4±0.5(bc) | 95.6±0.8(b) | 202.0±0.5(bc) | 104.1±0.8(b) |
| *Vrn-A1a+Vrn-D1a* | 8 | 186.9±0.3(c) | 92.1±0.4(c) | 200.6±0.2(c) | 100.6±0.4(c) |
| *vrn-B1+vrn-D1* | 35 | 192.8±0.4(a) | 98.4±0.4(a) | 204.5±0.3(a) | 106.1±0.4(a) |
| *vrn-B1+Vrn-D1a* | 19 | 189.2±0.2(b) | 94.5±0.2(b) | 202.1±0.1(b) | 102.7±0.3(b) |
| *Vrn-B1a+vrn-D1* | 3 | 188.0±0.8(bc) | 95.2±0.4(b) | 201.8±0.7(b) | 103.6±0.6(ab) |
| *Vrn-B1a+Vrn-D1a* | 4 | 186.4±0.6(c) | 91.0±0.3(c) | 200.4±0.3(b) | 100.1±0.3(b) |
| *Ppd-A1a+Ppd-B1a* | 4 | 187.7±0.2(c) | 93.0±0.5(b) | 201.2±0.2(c) | 100.9±0.4(b) |
| *Ppd-A1a+Ppd-B1b* | 4 | 192.5±1.2(ab) | 98.5±0.8(a) | 204.6±0.8(ab) | 106.5±0.6(a) |
| *Ppd-A1b+Ppd-B1a* | 37 | 190.3±0.3(bc) | 96.0±0.3(a) | 202.8±0.2(bc) | 104.1±0.3(a) |
| *Ppd-A1b+Ppd-B1b* | 16 | 193.2±0.6(a) | 98.2±0.7(a) | 204.7±0.4(a) | 106.1±0.7(a) |
| *Ppd-A1a+Ppd- D1b* | 7 | 201.9±0.2(a) | 104.5±0.1(a) | 211.3±0.0(a) | 111.5±0.2(a) |
| *Ppd-A1a+Ppd-D1a* | 1 | 188.4±0.3(c) | 94.5±0.4(b) | 201.7±0.2(c) | 102.6±0.4(b) |
| *Ppd-A1b+Ppd-D1b* | 12 | 196.7±0.7(b) | 103.1±0.6(a) | 207.8±0.5(b) | 110.8±0.6(a) |
| *Ppd-A1b+Ppd-D1a* | 41 | 189.5±0.2(c) | 94.8±0.2(b) | 202.1±0.1(c) | 102.9±0.2(b) |
| *Ppd-B1a +Ppd-D1b* | 6 | 195.2±0.9(b) | 101.9±0.6(b) | 206.9±0.7(b) | 109.6±0.4(b) |
| *Ppd-B1a +Ppd-D1a* | 35 | 189.1±0.2(c) | 94.7±0.2(c) | 201.9±0.1(c) | 102.7±0.2(c) |
| *Ppd-B1b+Ppd-D1b* | 7 | 198.7±0.9(a) | 104.3±0.9(a) | 209.2±0.6(a) | 111.9±0.9(a) |
| *Ppd-B1b+Ppd-D1a* | 13 | 190.1±0.2(c) | 95.0±0.3(c) | 202.3±0.1(c) | 103.1±0.3(c) |

**Note:** N: Number of varieties with the allelic combination. Different Letters in parentheses indicate a significant difference at the 0.05 level; decimal values preceded by “±” indicate standard error.

**Table 4**

The interactive effects of the allelic combinations of *VRN-1* and *PPD-1* loci on phenology stages in autumn sowing conditions (ASC) and spring sowing conditions (SSC)

| Allelic combinations | N | Days to heading | | Days to flowering | |
| --- | --- | --- | --- | --- | --- |
|  |  | **ASC** | **SSC** | **ASC** | **SSC** |
| *vrn-A1+Ppd-A1a* | 6 | 191.2±0.8(a) | 96.5±0.9(ab) | 203.6±0.6(a) | 104.3±0.9(ab) |
| *vrn-A1+Ppd-A1b* | 38 | 192.5±0.3(a) | 97.8±0.5(a) | 204.2±0.2(a) | 105.5±0.4(a) |
| *Vrn-A1a+Ppd-A1a* | 2 | 186.9±0.3(b) | 93.6±1.9(b) | 200.6±0.3(b) | 101.7±1.5(b) |
| *Vrn-A1a+Ppd-A1b* | 15 | 187.8±0.4(b) | 94.0±0.7(b) | 201.4±0.3(b) | 102.5±0.8(b) |
| *vrn-B1+Ppd-B1a* | 35 | 190.6±0.3(b) | 96.3±0.4(ab) | 203.1±0.2(a) | 104.2±0.4(ab) |
| *vrn-B1+Ppd-B1b* | 19 | 193.1±0.5(a) | 98.4±0.8(a) | 204.7±0.4(a) | 106.2±0.8(a) |
| *Vrn-B1a+Ppd-B1a* | 6 | 186.3±0.4(c) | 92.2±0.5(b) | 200.4±0.3(b) | 101.0±0.5(b) |
| *Vrn-B1a+Ppd-B1b* | 1 | 192.0±0.1(ab) | 96.4±0.3(ab) | 204.9±0.2(a) | 105.1±0.1(ab) |
| *vrn-D1+Ppd-D1b* | 12 | 197.8±0.7(a) | 103.6±0.9(a) | 208.5±0.5(a) | 111.2±0.8(a) |
| *vrn-D1+Ppd-D1a* | 26 | 189.9±0.3(b) | 95.7±0.4(b) | 202.3±0.2(b) | 103.5±0.4(b) |
| *Vrn-D1a+Ppd-D1a* | 22 | 188.7±0.2(c) | 93.7±0.3(b) | 201.7±0.1(b) | 102.0±0.3(b) |
| *Vrn-D1a+Ppd-D1b* | 1 | 189.0±0.1(bc) | 97.9±0.2(ab) | 202.9±0.1(b) | 106.6±0.1(ab) |

**Note:** N: Number of varieties with the allelic combination. Different Letters in parentheses indicate a significant

difference at the 0.05 level; decimal values preceded by “±” indicate standard error.

**Table 5**

The effects of two allelic combinations of *VRN-1*, *PPD-1* loci on morphological traits in autumn sowing conditions (ASC) and spring sowing conditions (SSC)

| Allelic combinations | N | Plant height (cm) | | Flag leaf length (cm) | | Flag leaf width (cm) | | Flag leaf area(cm^2^) | |
| --- | --- | --- | --- | --- | --- | --- | --- | --- | --- |
|  |  | **ASC** | **SSC** | **ASC** | **SSC** | **ASC** | **SSC** | **ASC** | **SSC** |
| *vrn-A1+vrn-B1* | 41 | 97.5±1.0(a) | 86.3±1.5(a) | 18.6 ± 0.3(a) | 22.5±0.4(a) | 1.43±0.01(a) | 1.67±0.01(a) | 29.6±1.2(a) | 37.5±0.6(a) |
| *vrn-A1+Vrn-B1a* | 3 | 94.0±1.6(ab) | 78.0±2.0(ab) | 15.9 ± 0.6(ab) | 20.2±0.4(ab) | 1.44±0.09(a) | 1.52±0.04(a) | 29.5±0.5(a) | 30.5±1.3(b) |
| *Vrn-A1a+vrn-B1* | 13 | 88.5±1.8(b) | 75.6±2.6(b) | 15.8 ± 0.4(b) | 20.1±0.3(b) | 1.43±0.03(a) | 1.67±0.03(a) | 24.5±0.5(b) | 33.3±0.5(b) |
| *Vrn-A1a+Vrn-B1a* | 4 | 85.3±1.3(b) | 76.9±2.3(ab) | 18.0 ±0.5(ab) | 20.9±1.2(ab) | 1.48±0.03(a) | 1.64±0.05(a) | 24.4±1.0(b) | 34.1±1.0(ab) |
| *vrn-A1+vrn-D1* | 29 | 98.2±1.3(a) | 88.5±1.8(a) | 19.2±0.5(a) | 23.2±0.4(a) | 1.42±0.02(a) | 1.70±0.02(a) | 30.1±0.7(a) | 39.0±0.7(a) |
| *vrn-A1+Vrn-D1a* | 15 | 95.3±1.2(ab) | 79.2±2.1(b) | 16.7±0.4(b) | 20.1±0.4(b) | 1.47±0.03(a) | 1.60±0.02(a) | 26.9±0.6(b) | 32.7±0.7(b) |
| *Vrn-A1a+vrn-D1* | 9 | 86.3±2.2(c) | 76.0±3.6(b) | 16.1±0.4(b) | 20.7±0.5(b) | 1.44±0.03(a) | 1.70±0.04(a) | 24.9±0.7(b) | 34.8±0.6(b) |
| *Vrn-A1a+Vrn-D1a* | 8 | 89.2±1.4(bc) | 75.8±1.4(b) | 16.7±0.5(b) | 19.9±0.4(b) | 1.44±0.03(a) | 1.62±0.03(a) | 26.9±0.8(b) | 32.2±0.6(b) |
| *vrn-B1+vrn-D1* | 35 | 96.4±1.2(a) | 86.8±1.8(a) | 18.7±0.4(a) | 22.7±0.4(a) | 1.42±0.01(a) | 1.68±0.02(a) | 29.2±0.6(a) | 38.2±0.6(a) |
| *vrn-B1+Vrn-D1a* | 19 | 93.5±1.2(ab) | 77.2±1.7(b) | 16.4±0.3(b) | 20.1±0.3(b) | 1.46±0.02(a) | 1.65±0.02(a) | 26.4±0.5(b) | 33.0±0.6(b) |
| *Vrn-B1a+vrn-D1* | 3 | 84.3±1.8(b) | 72.0±2.0(b) | 16.2±0.4(b) | 21.6±0.9(ab) | 1.46±0.05(a) | 1.70±0.09(a) | 25.9±1.3(b) | 36.3±1.0(ab) |
| *Vrn-B1a+Vrn-D1a* | 4 | 91.2±1.3(ab) | 80.5±1.8(ab) | 17.9±0.6(ab) | 20.0±0.6(b) | 1.46±0.06(a) | 1.53±0.02(b) | 28.7±1.2(ab) | 30.7±0.8(b) |
| *Ppd-A1a+Ppd-B1a* | 4 | 96.2±1.8(ab) | 83.7±1.0(ab) | 17.9±0.5(a) | 20.9±0.3(ab) | 1.40±0.04(b) | 1.51±0.03(c) | 27.9±0.9(a) | 31.5±0.6(c) |
| *Ppd-A1a+Ppd-B1b* | 4 | 101.8±2.1(a) | 87.5±3.1(ab) | 18.4±1.1(a) | 22.6±0.8(a) | 1.41±0.05(ab) | 1.63±0.02(bc) | 28.9±1.9(a) | 36.6±1.1(ab) |
| *Ppd-A1b+Ppd-B1a* | 37 | 91.9±1.2(b) | 78.0±1.1(b) | 17.1±0.4(a) | 20.6±0.2(b) | 1.49±0.02(a) | 1.71±0.01(a) | 28.0±0.6(a) | 35.0±0.4(b) |
| *Ppd-A1b+Ppd-B1b* | 16 | 99.1±1.6(a) | 88.1±2.1(a) | 18.4±0.4(a) | 22.7±0.4(a) | 1.41±0.03(b) | 1.63±0.02(b) | 28.0±0.6(a) | 37.0±0.7(a) |
| *Ppd-A1a+Ppd-D1a* | 7 | 97.1±1.4(ab) | 82.0±1.0(b) | 17.0±0.4(b) | 20.9±0.3(c) | 1.41±0.03(ab) | 1.57±0.02(b) | 26.7±0.7(b) | 32.7±0.5(b) |
| *Ppd-A1a+Ppd-D1b* | 1 | 112.5±1.9(a) | 111.1±0.7(a) | 26.1±2.2(a) | 28.1±2.0(a) | 1.37±0.12(ab) | 1.55±0.04(b) | 40.1±4.8(a) | 43.7±2.3(a) |
| *Ppd-A1b+Ppd-D1b* | 12 | 104.1±1.9(a) | 97.4±2.3(a) | 21.1±0.6(a) | 24.6±0.4(b) | 1.35±0.02(b) | 1.61±0.02(b) | 31.7±1.0(a) | 39.7±0.8(a) |
| *Ppd-A1b+Ppd-D1a* | 41 | 91.1±1.1(b) | 76.2±0.9(b) | 16.4±0.3(b) | 20.2±0.2(c) | 1.50±0.02(a) | 1.71±0.01(a) | 26.9±0.5(b) | 34.4±0.3(b) |
| *Ppd-B1a +Ppd-D1b* | 6 | 101.8±2.4(ab) | 94.6±3.3(a) | 21.6±0.9(a) | 24.4±0.6(a) | 1.42±0.03(ab) | 1.71±0.02(a) | 34.5±1.7(a) | 41.6±1.1(a) |
| *Ppd-B1a +Ppd-D1a* | 35 | 90.7±1.1(c) | 75.8±0.9(b) | 16.4±0.4(b) | 19.9±0.2(c) | 1.49±0.02(a) | 1.68±0.01(a) | 26.8±0.6(c) | 33.5±0.3(c) |
| *Ppd-B1b+Ppd-D1b* | 7 | 107.4±2.5(a) | 101.8±2.6(a) | 21.4±0.8(a) | 25.3±0.6(a) | 1.30±0.03(b) | 1.52±0.01(b) | 30.6±1.2(ab) | 38.6±1.0(ab) |
| *Ppd-B1b+Ppd-D1a* | 13 | 95.5±1.5(bc) | 80.5±1.8(b) | 16.8±0.4(b) | 21.3±0.3(b) | 1.47±0.03(a) | 1.69±0.02(a) | 26.9±0.6(bc) | 36.0±0.7(b) |

**Note:** N: Number of varieties with the allelic combination. Different Letters in parentheses indicate a significant difference at the 0.05 level; decimal values preceded by “±” indicate standard error.

**Table 6**

The interactive effects of two allelic combinations of *PPD-1 and VRN-1* loci on morphological traits in autumn sowing conditions (ASC) and spring sowing conditions (SSC)

| Allelic combinations | N | Plant height (cm) | | Flag leaf length (cm) | | Flag leaf width (cm) | | Flag leaf area(cm^2^) | |
| --- | --- | --- | --- | --- | --- | --- | --- | --- | --- |
|  |  | **ASC** | **SSC** | **ASC** | **SSC** | **ASC** | **SSC** | **ASC** | **SSC** |
| *vrn-A1+Ppd-A1a* | 6 | 100.9±1.7(a) | 86.9±2.9(a) | 18.3±0.8(a) | 22.3±0.8(a) | 1.38±0.03(a) | 1.54±0.03(b) | 27.9±1.2(ab) | 34.4±1.3(a) |
| *vrn-A1+Ppd-A1b* | 38 | 96.6±1.1(a) | 83.5±1.7(a) | 18.0±0.4(a) | 21.6±0.3(a) | 1.47±0.02(a) | 1.68±0.02(a) | 29.0±0.6(a) | 36.4±0.6(a) |
| *Vrn-A1a+Ppd-A1a* | 2 | 93.3±2.2(ab) | 81.9±3.7(ab) | 17.8±1.0(ab) | 20.1±0.7(a) | 1.50±0.05(a) | 1.64±0.07(ab) | 29.9±1.7(a) | 33.1±1.5(a) |
| *Vrn-A1a+Ppd-A1b* | 15 | 87.6±1.7(b) | 74.8±2.5(ab) | 16.1±0.4(ab) | 20.2±0.4(a) | 1.45±0.02(a) | 1.68±0.03(a) | 25.3±0.6(ab) | 33.8±0.5(a) |
| *vrn-B1+Ppd-B1a* | 35 | 92.9±1.2(b) | 78.8±1.6(b) | 17.1±0.4(a) | 20.7±0.3(ab) | 1.47±0.02(a) | 1.69±0.02(a) | 27.8±0.7(a) | 35.0±0.5(ab) |
| *vrn-B1+Ppd-B1b* | 19 | 100.1±1.4(a) | 88.4±2.6(a) | 18.5±0.4(a) | 22.6±0.5(a) | 1.42±0.02(a) | 1.64±0.02(ab) | 28.4±0.6(a) | 37.1±0.8(a) |
| *Vrn-B1a+Ppd-B1a* | 6 | 88.8±1.4(b) | 77.0±2.1(b) | 17.5±0.5(a) | 19.8±0.5(ab) | 1.52±0.05(a) | 1.65±0.04(ab) | 29.1±1.1(a) | 32.6±1.1(b) |
| *Vrn-B1a+Ppd-B1b* | 1 | 91.2±3.1(b) | 79.9±0.5(ab) | 16.5±0.8(a) | 24.4±0.5(a) | 1.26±0.08(a) | 1.40±0.01(b) | 23.0±1.3(a) | 34.3±0.5(ab) |
| *vrn-D1+Ppd-D1b* | 12 | 106.1±1.8(a) | 99.9±3.1(a) | 21.9±0.6(a) | 25.5±0.5(a) | 1.35±0.03(b) | 1.61±0.02(b) | 32.9±1.1(a) | 41.1±0.9(a) |
| *vrn-D1+Ppd-D1a* | 26 | 90.7±1.5(b) | 76.4±1.7(b) | 16.3±0.5(b) | 20.5±0.3(b) | 1.49±0.02(a) | 1.73±0.02(a) | 26.6±0.7(b) | 35.4±0.6(b) |
| *Vrn-D1a+Ppd-D1a* | 22 | 89.1±1.1(b) | 81.0±3.9(ab) | 16.8±0.6(ab) | 17.5±0.4(b) | 1.40±0.08(ab) | 1.51±0.03(b) | 26.1±1.0(b) | 26.6±1.1(c) |
| *Vrn-D1a+Ppd-D1b* | 1 | 93.6±1.0(b) | 77.8±1.5(b) | 16.7±0.3(b) | 20.1±0.3(b) | 1.47±0.02(a) | 1.64±0.02(b) | 27.1±0.5(b) | 32.8±0.5(c) |

**Note:** N: Number of varieties with the allelic combination. Different Letters in parentheses indicate a significant difference at the 0.05 level; decimal values preceded by “±” indicate standard error.

**Table 7**

The effects of two allelic combinations of *VRN-1*, *PPD-1* loci on spike traits in autumn sowing conditions (ASC) and spring sowing conditions (SSC)

| Allelic combinations | N | Grain number per spike | | Spike length (cm) | | Spikelet per spike | | Spike Fertility  (GN g chaff^-1^) | |
| --- | --- | --- | --- | --- | --- | --- | --- | --- | --- |
|  |  | **ASC** | **SSC** | **ASC** | **SSC** | **ASC** | **SSC** | **ASC** | **SSC** |
| *vrn-A1+vrn-B1* | 41 | 42.8±0.4(a) | 47.8±0.5(a) | 9.6±0.1(b) | 9.2±0.1(a) | 19.0±0.1(a) | 19.5±0.1(a) | 77.4±NA(a) | 76.8±1.5(a) |
| *vrn-A1+Vrn-B1a* | 3 | 45.6±1.7(a) | 43.5±0.9(ab) | 10.2±0.1(ab) | 8.9±0.1(a) | 17.8±0.2(b) | 17.4±0.3(c) | 76.9±2.2(a) | 75.4±5.8(a) |
| *Vrn-A1a+vrn-B1* | 13 | 43.1±0.8(a) | 46.3±0.7(ab) | 9.4±0.1(b) | 8.8±0.2(a) | 19.0±0.2(a) | 18.9±0.2(b) | 71.2±2.8(b) | 82.6±2.3(a) |
| *Vrn-A1a+Vrn-B1a* | 4 | 43.6±1.5(a) | 47.9±2.1(a) | 10.7±0.2(a) | 9.0±0.1(a) | 17.1±0.2(b) | 17.2±0.6(c) | 65.8±2.9(b) | 82.2±3.0(a) |
| *vrn-A1+vrn-D1* | 29 | 43.0±0.5(a) | 47.8±0.6(a) | 9.7±0.1(a) | 9.5±0.2(a) | 19.1±0.1(a) | 19.6±0.1(a) | 76.1±NA(ab) | 74.1±1.7(ab) |
| *vrn-A1+Vrn-D1a* | 15 | 43.0±0.6(a) | 46.8±0.5(ab) | 9.3±0.1(a) | 8.5±0.1(b) | 18.4±0.1(b) | 18.8±0.1(b) | 79.7±2.2(a) | 82.0±2.3(a) |
| *Vrn-A1a+vrn-D1* | 9 | 42.8±1.0(a) | 49.1±1.0(a) | 9.7±0.2(a) | 9.2±0.2(ab) | 19.3±0.2(a) | 19.4±0.3(ab) | 69.7±3.1(b) | 82.3±2.5(a) |
| *Vrn-A1a+Vrn-D1a* | 8 | 43.7±0.9(a) | 44.3±1.0(b) | 9.7±0.2(a) | 8.6±0.2(b) | 17.7±0.2(c) | 17.5±0.3(c) | 70.2±3.4(ab) | 82.6±2.8(a) |
| *vrn-B1+vrn-D1* | 35 | 43.2±0.5(b) | 47.8±0.5(ab) | 9.7±0.1(b) | 9.5±0.1(a) | 19.3±0.1(a) | 19.7±0.1(a) | 74.8±NA(a) | 75.5±1.6(b) |
| *vrn-B1+Vrn-D1a* | 19 | 42.3±0.5(b) | 46.7±0.5(b) | 9.2±0.1(c) | 8.5±0.1(b) | 18.4±0.1(b) | 18.7±0.1(b) | 77.7±2.2(a) | 83.3±2.0(a) |
| *Vrn-B1a+vrn-D1* | 3 | 40.0±1.6(b) | 51.1±2.1(a) | 10.3±0.2(ab) | 9.2±0.1(ab) | 17.5±0.2(bc) | 18.1±0.5(b) | 70.9±2.6(a) | 81.2±4.8(ab) |
| *Vrn-B1a+Vrn-D1a* | 4 | 47.8±1.2(a) | 42.2±1.1(c) | 10.6±0.1(a) | 8.7±0.1(ab) | 17.3±0.3(c) | 16.6±0.4(c) | 70.3±3.1(a) | 77.8±3.9(ab) |
| *Ppd-A1a+Ppd-B1a* | 4 | 43.0±0.9(a) | 48.0±1.2(a) | 10.0±0.2(a) | 8.6±0.1(b) | 17.8±0.3(b) | 17.3±0.5(b) | 67.8±2.3(b) | 79.4±2.8(a) |
| *Ppd-A1a+Ppd-B1b* | 4 | 41.9±0.8(a) | 44.1±1.3(a) | 10.0±0.2(a) | 10.9±0.3(a) | 18.8±0.2(ab) | 19.6±0.3(a) | 74.9±6.8(ab) | 79.3±4.6(a) |
| *Ppd-A1b+Ppd-B1a* | 37 | 43.8±0.5(a) | 47.8±0.6(a) | 9.6±0.1(a) | 9.0±0.1(b) | 18.8±0.1(a) | 19.1±0.1(a) | 74.0±NA(b) | 78.2±1.4(a) |
| *Ppd-A1b+Ppd-B1b* | 16 | 41.7±0.6(a) | 46.5±0.6(a) | 9.5±0.1(a) | 9.1±0.2(b) | 19.0±0.1(a) | 19.4±0.1(a) | 80.0±2.2(a) | 78.4±2.8(a) |
| *Ppd-A1a+Ppd-D1b* | 7 | 42.7±0.7(a) | 46.0±1.0(b) | 9.8±0.1(b) | 9.2±0.2(c) | 18.1±0.2(ab) | 18.3±0.3(b) | 76.2±2.7(a) | 82.8±2.5(a) |
| *Ppd-A1a+Ppd-D1a* | 1 | 40.8±0.2(a) | 46.3±2.2(b) | 11.0±0.2(a) | 12.2±0.9(a) | 19.7±0.6(a) | 19.7±0.6(a) | 37.5±16.8(b) | 55.1±3.9(b) |
| *Ppd-A1b+Ppd-D1b* | 12 | 42.5±0.7(a) | 49.8±1.0(a) | 10.7±0.2(a) | 10.4±0.3(b) | 19.4±0.2(a) | 20.2±0.2(a) | 77.8±NA(a) | 73.9±3.2(ab) |
| *Ppd-A1b+Ppd-D1a* | 41 | 43.3±0.5(a) | 46.7±0.5(b) | 9.2±0.1(c) | 8.6±0.1(c) | 18.7±0.1(ab) | 18.9±0.1(b) | 75.3±1.4(a) | 79.6±1.3(a) |
| *Ppd-B1a +Ppd-D1b* | 6 | 44.6±1.1(a) | 54.3±1.3(a) | 10.9±0.3(a) | 10.9±0.4(a) | 19.5±0.3(a) | 20.8±0.1(a) | 77.1±NA(ab) | 69.2±1.6(b) |
| *Ppd-B1a +Ppd-D1a* | 35 | 43.6±0.5(a) | 46.3±0.5(b) | 9.4±0.1(b) | 8.6±0.1(b) | 18.6±0.1(b) | 18.6±0.1(c) | 72.8±1.4(b) | 79.8±1.4(a) |
| *Ppd-B1b+Ppd-D1b* | 7 | 40.6±0.8(b) | 46.1±0.8(b) | 10.6±0.2(a) | 10.3±0.3(a) | 19.3±0.2(a) | 19.5±0.2(b) | 72.6±3.8(b) | 74.6±5.0(ab) |
| *Ppd-B1b+Ppd-D1a* | 13 | 42.3±0.7(ab) | 47.3±0.7(b) | 9.1±0.1(b) | 8.8±0.2(b) | 18.8±0.1(ab) | 19.3±0.1(b) | 82.4±2.6(a) | 80.9±2.3(a) |

**Note:** N: Number of varieties with the allelic combination. Different Letters in parentheses indicate a significant difference at the 0.05 level; decimal values preceded by “±” indicate standard error.

**Table 8**

The effects of two allelic combinations of *VRN-1*, *PPD-1* loci on yield traits in autumn sowing conditions (ASC) and spring sowing conditions (SSC)

| Allelic combinations | N | Thousand kernel weight (g) | | Grain yield (gm^-2^) | | Biomass (gm^-2^) | | Harvest Index | |
| --- | --- | --- | --- | --- | --- | --- | --- | --- | --- |
|  |  | **ASC** | **SSC** | **ASC** | **SSC** | **ASC** | **SSC** | **ASC** | **SSC** |
| *vrn-A1+vrn-B1* | 41 | 45.6±0.5(ab) | 37.5±0.3(b) | 430.5±8.4(a) | 286.8±5.8(b) | 1049.5±15.2(a) | 730.3±12.2(a) | 0.42±0.01(ab) | 0.40±0.01(b) |
| *vrn-A1+Vrn-B1a* | 3 | 42.4±1.9(b) | 37.2±0.9(b) | 397.0±14.9(a) | 335.2±24.3(ab) | 1083.9±30.5(a) | 789.4±49.7(a) | 0.37±0.02(b) | 0.43±0.02(ab) |
| *Vrn-A1a+vrn-B1* | 13 | 46.3±0.8(a) | 37.5±0.4(b) | 438.9±15.9(a) | 314.7±10.9(b) | 1076.0±32.1(a) | 719.7±21.1(a) | 0.42±0.02(ab) | 0.45±0.02(a) |
| *Vrn-A1a+Vrn-B1a* | 4 | 43.7±1.8(ab) | 41.3±0.8(a) | 418.2±20.4(a) | 390.4±12.9(a) | 948.1±31.5(ab) | 816.6±17.4(a) | 0.45±0.02(a) | 0.48±0.01(a) |
| *vrn-A1+vrn-D1* | 29 | 44.9±0.5(a) | 37.7±0.4(ab) | 427.0±10.1(a) | 277.9±6.9(b) | 1051.7±18.5(a) | 735.0±14.8(a) | 0.41±0.01(a) | 0.38±0.01(b) |
| *vrn-A1+Vrn-D1a* | 15 | 45.7±0.9(a) | 36.9±0.3(b) | 430.8±12.4(a) | 316.5±9.8(a) | 1039.2±22.3(a) | 734.1±20.1(a) | 0.42±0.01(a) | 0.44±0.01(a) |
| *Vrn-A1a+vrn-D1* | 9 | 46.3±1.1(a) | 39.4±0.7(a) | 440.0±15.4(a) | 329.1±12.6(a) | 1063.5±37.6(a) | 759.5±19.9(a) | 0.43±0.01(a) | 0.43±0.01(a) |
| *Vrn-A1a+Vrn-D1a* | 8 | 45.0±1.1(a) | 37.4±0.5(ab) | 427.3±21.7(a) | 336.3±14.2(a) | 1026.1±36.1(a) | 723.3±28.6(a) | 0.42±0.02(a) | 0.48±0.02(a) |
| *vrn-B1+vrn-D1* | 35 | 45.2±0.5(ab) | 37.9±0.3(ab) | 429.9±9.1(a) | 284.1±6.8(c) | 1053.5±17.9(a) | 731.1±13.6(b) | 0.41±0.01(a) | 0.39±0.01(b) |
| *vrn-B1+Vrn-D1a* | 19 | 46.4±0.7(a) | 36.7±0.3(b) | 437.5±12.7(a) | 313.4±8.7(b) | 1050.1±21.9(a) | 720.7±19.4(b) | 0.42±0.01(a) | 0.45±0.01(a) |
| *Vrn-B1a+vrn-D1* | 3 | 46.0±1.8(ab) | 40.5±1.0(a) | 432.2±19.5(a) | 363.3±14.7(ab) | 1066.5±30.8(a) | 852.1±22.5(a) | 0.42±0.03(a) | 0.43±0.01(ab) |
| *Vrn-B1a+Vrn-D1a* | 4 | 41.0±1.7(b) | 38.9±0.9(ab) | 391.7±17.5(a) | 369.3±20.7(a) | 961.1±33.5(a) | 769.5±35.7(ab) | 0.41±0.02(a) | 0.48±0.02(a) |
| *Ppd-A1a+Ppd-B1a* | 4 | 47.2±1.7(a) | 40.7±0.5(a) | 419.3±24.6(a) | 378.1±10.3(a) | 1051.4±47.4(a) | 826.2±23.5(a) | 0.40±0.03(a) | 0.46±0.01(a) |
| *Ppd-A1a+Ppd-B1b* | 4 | 46.2±1.1(a) | 37.1±0.7(bc) | 485.2±19.4(a) | 272.9±12.8(c) | 1124.7±36.4(a) | 751.2±54.2(a) | 0.43±0.02(a) | 0.39±0.03(ab) |
| *Ppd-A1b+Ppd-B1a* | 37 | 46.0±0.4(a) | 38.0±0.3(b) | 432.2±8.5(a) | 310.5±6.2(b) | 1031.7±16.5(a) | 731.8±13.3(a) | 0.43±0.01(a) | 0.43±0.01(a) |
| *Ppd-A1b+Ppd-B1b* | 16 | 43.2±0.9(ab) | 36.4±0.5(c) | 413.2±14.2(a) | 273.9±11.2(c) | 1061.9±25.1(a) | 721.3±19.7(a) | 0.39±0.01(a) | 0.38±0.01(b) |
| *Ppd-A1a+Ppd-D1a* | 7 | 46.9±1.1(a) | 38.4±0.5(a) | 433.9±16.6(bc) | 331.9±12.9(a) | 1053.5±30.7(bc) | 759.9±26.6(b) | 0.42±0.02(a) | 0.45±0.02(a) |
| *Ppd-A1a+Ppd-D1b* | 1 | 45.45±0.1(ab) | 42.7±0.49(a) | 580.6±15.3(a) | 280.7±35.7(a) | 1329.8±16.1(a) | 990.3±60.8(a) | 0.44±0.01(a) | 0.29±0.06(b) |
| *Ppd-A1b+Ppd-D1b* | 12 | 41.8±1.0(b) | 35.4±0.5(b) | 389.0±14.7(c) | 249.6±10.9(b) | 1125.9±30.0(ab) | 770.6±22.2(b) | 0.35±0.01(ab) | 0.32±0.01(b) |
| *Ppd-A1b+Ppd-D1a* | 41 | 46.1±0.4(a) | 38.2±0.3(a) | 437.4±8.4(b) | 315.5±6.5(a) | 1015.9±15.2(c) | 715.0±12.6(b) | 0.43±0.01(a) | 0.44±0.01(a) |
| *Ppd-B1a +Ppd-D1b* | 6 | 43.8±1.3(ab) | 36.3±0.8(b) | 367.3±20.8(b) | 287.7±9.7(b) | 1080.5±42.0(ab) | 825.3±19.1(a) | 0.35±0.02(c) | 0.34±0.01(b) |
| *Ppd-B1a +Ppd-D1a* | 35 | 46.5±0.4(a) | 38.7±0.3(a) | 441.9±8.5(a) | 323.3±6.6(a) | 1025.6±16.7(b) | 726.1±13(b) | 0.44±0.01(a) | 0.45±0.01(a) |
| *Ppd-B1b+Ppd-D1b* | 7 | 40.6±1.2(b) | 35.7±0.7(b) | 435.0±19.8(ab) | 221.3±11.8(b) | 1193.9±37.1(a) | 755.1±29(ab) | 0.37±0.02(bc) | 0.30±0.01(b) |
| *Ppd-B1b+Ppd-D1a* | 13 | 45.5±1.0(a) | 37.0±0.5(b) | 423.6±15.7(ab) | 304.3±12.5(a) | 1010.1±23.3(b) | 711.6±24.3(b) | 0.42±0.01(ab) | 0.43±0.01(a) |

**Note:** N: Number of varieties with the allelic combination. Different Letters in parentheses indicate a significant difference at the 0.05 level; decimal values preceded by “±” indicate standard error.

**Table 9**

The interactive effects of two allelic combinations of *VRN-1* with *PPD-1* on spike and yield traits in autumn sowing conditions (ASC) and spring sowing conditions (SSC)

| Allelic combinations | N | Grain number per spike | | Spike length (cm) | | Spikelet per spike | | Spike Fertility  (GN g chaff^-1^) | |
| --- | --- | --- | --- | --- | --- | --- | --- | --- | --- |
|  |  | **ASC** | **SSC** | **ASC** | **SSC** | **ASC** | **SSC** | **ASC** | **SSC** |
| *vrn-A1+Ppd-A1a* | 6 | 42.6±0.7(a) | 46.7±1.2(a) | 9.8±0.1(ab) | 9.8±0.3(a) | 18.7±0.2(a) | 19.0±0.3(a) | 70.9±4.1(ab) | 75.8±3.2(b) |
| *vrn-A1+Ppd-A1b* | 38 | 43.0±0.5(a) | 41.3±1.6(a) | 9.6±0.1(b) | 9.1±0.1(ab) | 18.9±0.1(a) | 19.4±0.1(a) | 78.4±NA(a) | 76.1±1.6(b) |
| *Vrn-A1a+Ppd-A1a* | 2 | 42.1±1.4(a) | 44.2±4.0(a) | 10.6±0.2(a) | 9.1±0.3(ab) | 17.0±0.5(b) | 16.7±1.5(b) | 72.7±7.8(ab) | 76.6±3.8(ab) |
| *Vrn-A1a+Ppd-A1b* | 15 | 43.4±0.8(a) | 43.9±2.0(a) | 9.6±0.1(ab) | 8.8±0.2(b) | 18.7±0.2(a) | 18.8±0.3(a) | 69.5±2.4(b) | 84.1±2.0(a) |
| *vrn-B1+Ppd-B1a* | 35 | 43.3±0.5(ab) | 40.8±1.7(a) | 9.5±0.1(b) | 7.2±0.4(a) | 18.9±0.1(a) | 19.2±0.2(a) | 73.9±NA(a) | 77.3±3.1(ab) |
| *vrn-B1+Ppd-B1b* | 19 | 42.0±0.5(b) | 44.6±1.6(a) | 9.6±0.1(b) | 8.4±0.4(a) | 19.0±0.1(a) | 19.6±0.2(a) | 79.4±2.3(a) | 79.8±4.0(a) |
| *Vrn-B1a+Ppd-B1a* | 6 | 45.6±1.2(a) | 46.4±2.2(a) | 10.5±0.1(a) | 9.0±0.2(a) | 17.4±0.2(b) | 17.4±0.5(b) | 70.5±2.4(a) | 83.2±4.4(a) |
| *Vrn-B1a+Ppd-B1b* | 1 | 37.2±0.8(b) | 43.5±1.6(a) | 10.2±0.1(ab) | 8.7±0.3(a) | 17.7±0.2(ab) | 16.4±0.2(b) | 71.1±1.0(a) | 56.0±1.9(a) |
| *vrn-D1+Ppd-D1b* | 12 | 42.3±0.7(a) | 50.1±2.6(a) | 10.9±0.2(a) | 9.9±0.6(a) | 19.4±0.2(a) | 20.2±0.3(a) | 74.4±NA(a) | 72.5±5.4(b) |
| *vrn-D1+Ppd-D1a* | 26 | 43.2±0.6(a) | 41.7±1.9(b) | 9.2±0.1(b) | 7.1±0.4(b) | 19.0±0.1(a) | 19.3±0.2(b) | 74.6±1.6(a) | 77.7±3.3(ab) |
| *Vrn-D1a+Ppd-D1a* | 22 | 42.4±1.0(a) | 42.9±1.1(a) | 9.4±0.1(b) | 8.3±0.3(ab) | 18.7±0.4(ab) | 19.6±0.4(ab) | 77.4±3.4(a) | 70.8±8.8(a) |
| *Vrn-D1a+Ppd-D1b* | 1 | 43.3±0.5(a) | 46.0±1.8(b) | 9.5±0.1(b) | 7.4±0.4(b) | 18.2±0.1(b) | 18.3±0.2(b) | 76.4±2.0(a) | 82.8±3.7(a) |
| Allelic combinations | **N** | **Thousand kernel weight (g)** | | **Grain yield (gm^-2^)** | | **Biomass (gm^-2^)** | | **Harvest Index** | |
|  |  | **ASC** | **SSC** | **ASC** | **SSC** | **ASC** | **SSC** | **ASC** | **SSC** |
| *vrn-A1+Ppd-A1a* | 6 | 47.3±0.8(a) | 39.0±0.6(a) | 474.8±15.8(a) | 322.0±21.4(ab) | 1130.1±31.6(a) | 803.7±49.2(a) | 0.43±0.02(a) | 0.42±0.03(ab) |
| *vrn-A1+Ppd-A1b* | 38 | 44.9±0.5(a) | 34.3±1.0(a) | 420.9±8.7(a) | 254.9±11.3(ab) | 1034.4±15.7(a) | 646.5±25.9(a) | 0.41±0.01(a) | 0.35±0.01(b) |
| *Vrn-A1a+Ppd-A1a* | 2 | 45.0±3.1(a) | 38.6±2.4(a) | 384.6±39.3(a) | 336.2±23.2(a) | 961.9±62.6(a) | 743.5±20.3(a) | 0.40±0.03(a) | 0.45±0.03(ab) |
| *Vrn-A1a+Ppd-A1b* | 15 | 45.8±0.8(a) | 35.8±1.6(a) | 440.6±13.8(a) | 332.0±14.8(a) | 1057.1±28.3(a) | 742.4±27.5(a) | 0.43±0.01(a) | 0.46±0.02(a) |
| *vrn-B1+Ppd-B1a* | 35 | 46.4±0.4(a) | 38.0±1.1(b) | 432.6±9.1(a) | 281.4±11.4(b) | 1042.8±17.7(a) | 733.5±26(ab) | 0.42±0.01(a) | 0.39±0.01(ab) |
| *vrn-B1+Ppd-B1b* | 19 | 44.2±0.8(ab) | 36.6±1.2(c) | 432.5±12.7(a) | 255.4±15.0(b) | 1069.7±22.4(a) | 716.9±32.2(b) | 0.41±0.01(ab) | 0.36±0.02(b) |
| *Vrn-B1a+Ppd-B1a* | 6 | 44.3±1.4(ab) | 40.3±1.0(a) | 421.5±14.2(a) | 369.9±21.8(a) | 979.9±24.9(a) | 785.6±36.8(a) | 0.43±0.01(a) | 0.47±0.02(a) |
| *Vrn-B1a+Ppd-B1b* | 1 | 36.3±0.1(b) | 35.2±0.4(a) | 334.8±19.6(a) | 347.7±30.0(ab) | 1164.4±47.2(a) | 920.8±24.5(a) | 0.30±0.03(b) | 0.38±0.04(ab) |
| *vrn-D1+Ppd-D1b* | 12 | 41.7±1.0(ab) | 35.9±0.8(b) | 401.8±15.8(a) | 253.8±15.6(b) | 1150.1±29.9(a) | 792.6±32.9(a) | 0.36±0.01(b) | 0.32±0.01(b) |
| *vrn-D1+Ppd-D1a* | 26 | 46.9±0.5(a) | 39.2±1.5(a) | 443.1±10.0(a) | 309.2±13.2(a) | 1010.4±19.1(b) | 715.4±27.6(b) | 0.44±0.01(a) | 0.40±0.02(a) |
| *Vrn-D1a+Ppd-D1a* | 22 | 47.0±0.5(a) | 36.8±1.1(b) | 427.0±35.6(a) | 230.5±29.8(a) | 1038.8±80.1(ab) | 726.0±77.5(a) | 0.41±0.03(ab) | 0.33±0.07(ab) |
| *Vrn-D1a+Ppd-D1b* | 1 | 45.4±0.7(a) | 37.1±1.1(b) | 429.7±11.4(a) | 328.8±15.8(a) | 1034.4±19.7(b) | 730.2±34.2(ab) | 0.42±0.01(a) | 0.42±0.02(a) |

**Note:** N: Number of varieties with the allelic combination. Different Letters in parentheses indicate a significant difference at the 0.05 level; decimal values preceded by “±” indicate standard error.

**Supplementary Table 1**

The origination based on the geographic region of wheat varieties and their allelic variants on *VRN-1* and *PPD-1* alleles

| s/n | Varieties | Origin | VRN-A1 | VRN-B1 | VRN-D1 | PPD-A1 | PPD-B1 | PPD-D1 |
| --- | --- | --- | --- | --- | --- | --- | --- | --- |
| 1 | Fengchan3 | SHWW | vrn-A1 | vrn-B1 | Vrn-D1a | Ppd-A1b | Ppd-B1a | Ppd-D1a |
| 2 | Jing411 | NWW | vrn-A1 | vrn-B1 | vrn-D1 | Ppd-A1b | Ppd-B1b | Ppd-D1a |
| 3 | Ningdong10 | NWWW | vrn-A1 | vrn-B1 | vrn-D1 | Ppd-A1b | Ppd-B1a | Ppd-D1b |
| 4 | Ningdong11 | NWWW | vrn-A1 | vrn-B1 | vrn-D1 | Ppd-A1b | Ppd-B1a | Ppd-D1a |
| 5 | Shan229 | SHWW | vrn-A1 | vrn-B1 | vrn-D1 | Ppd-A1a | Ppd-B1b | Ppd-D1a |
| 6 | Xifeng20 | NWW | vrn-A1 | vrn-B1 | Vrn-D1a | Ppd-A1b | Ppd-B1a | Ppd-D1a |
| 7 | Xiaoyan6 | SHWW | vrn-A1 | vrn-B1 | vrn-D1 | Ppd-A1b | Ppd-B1b | Ppd-D1a |
| 8 | Chang6878 | NWW | vrn-A1 | vrn-B1 | vrn-D1 | Ppd-A1a | Ppd-B1b | Ppd-D1a |
| 9 | Zhongmai175 | NWW | vrn-A1 | vrn-B1 | vrn-D1 | Ppd-A1b | Ppd-B1a | Ppd-D1a |
| 10 | Aifeng3 | SHWW | vrn-A1 | vrn-B1 | Vrn-D1a | Ppd-A1b | Ppd-B1b | Ppd-D1a |
| 11 | Bainong207 | SHWW | vrn-A1 | vrn-B1 | vrn-D1 | Ppd-A1b | Ppd-B1a | Ppd-D1a |
| 12 | FengDeCun1 | NHWW | vrn-A1 | vrn-B1 | vrn-D1 | Ppd-A1b | Ppd-B1a | Ppd-D1a |
| 13 | Huai Mai25 | SHWW | vrn-A1 | Vrn-B1a | Vrn-D1a | Ppd-A1b | Ppd-B1a | Ppd-D1a |
| 14 | Jining13 | NHWW | vrn-A1 | vrn-B1 | vrn-D1 | Ppd-A1b | Ppd-B1a | Ppd-D1b |
| 15 | Luomai22 | SHWW | vrn-A1 | vrn-B1 | vrn-D1 | Ppd-A1b | Ppd-B1a | Ppd-D1a |
| 16 | Xinong389 | SHWW | vrn-A1 | vrn-B1 | Vrn-D1a | Ppd-A1b | Ppd-B1a | Ppd-D1a |
| 17 | Xinong529 | SHWW | Vrn-A1a | Vrn-B1a | vrn-D1 | Ppd-A1b | Ppd-B1a | Ppd-D1a |
| 18 | Xinong979 | SHWW | Vrn-A1a | vrn-B1 | vrn-D1 | Ppd-A1b | Ppd-B1b | Ppd-D1a |
| 19 | Xinmai18 | NHWW | Vrn-A1a | vrn-B1 | vrn-D1 | Ppd-A1b | Ppd-B1a | Ppd-D1a |
| 20 | Xinmai26 | NHWW | Vrn-A1a | Vrn-B1a | vrn-D1 | Ppd-A1b | Ppd-B1a | Ppd-D1a |
| 21 | Changhan58 | NWW | vrn-A1 | vrn-B1 | vrn-D1 | Ppd-A1b | Ppd-B1b | Ppd-D1a |
| 22 | Zhengmai366 | SHWW | Vrn-A1a | vrn-B1 | vrn-D1 | Ppd-A1b | Ppd-B1a | Ppd-D1b |
| 23 | Zhoumai 18 | SHWW | Vrn-A1a | vrn-B1 | Vrn-D1a | Ppd-A1b | Ppd-B1a | Ppd-D1a |
| 24 | Clear white | Western USA | Vrn-A1a | Vrn-B1a | Vrn-D1a | Ppd-A1b | Ppd-B1a | Ppd-D1a |
| 25 | Lassit | Western USA | vrn-A1 | vrn-B1 | vrn-D1 | Ppd-A1b | Ppd-B1a | Ppd-D1a |
| 26 | Han6172 | NWW | vrn-A1 | vrn-B1 | Vrn-D1a | Ppd-A1a | Ppd-B1a | Ppd-D1a |
| 27 | Jinmai47 | NWW | vrn-A1 | vrn-B1 | Vrn-D1a | Ppd-A1a | Ppd-B1a | Ppd-D1a |
| 28 | Luohan2 | HHWW | vrn-A1 | vrn-B1 | Vrn-D1a | Ppd-A1b | Ppd-B1b | Ppd-D1b |
| 29 | Shijiazhuang8 | NWW | vrn-A1 | vrn-B1 | Vrn-D1a | Ppd-A1b | Ppd-B1a | Ppd-D1a |
| 30 | Taishan5 | NHWW | vrn-A1 | vrn-B1 | vrn-D1 | Ppd-A1b | Ppd-B1a | Ppd-D1a |
| 31 | Xinong928 | NWW | vrn-A1 | vrn-B1 | Vrn-D1a | Ppd-A1a | Ppd-B1a | Ppd-D1a |
| 32 | Yunhan618 | NWW | Vrn-A1a | vrn-B1 | vrn-D1 | Ppd-A1b | Ppd-B1a | Ppd-D1a |
| 33 | Zhangwu134 | NWW | vrn-A1 | vrn-B1 | Vrn-D1a | Ppd-A1b | Ppd-B1a | Ppd-D1a |
| 34 | Xinong511 | HHWW | Vrn-A1a | vrn-B1 | Vrn-D1a | Ppd-A1b | Ppd-B1b | Ppd-D1a |
| 35 | Drysdale | Australia | vrn-A1 | vrn-B1 | vrn-D1 | Ppd-A1b | Ppd-B1b | Ppd-D1a |
| 36 | Quarrion | Australia | vrn-A1 | Vrn-B1a | vrn-D1 | Ppd-A1b | Ppd-B1b | Ppd-D1b |
| 37 | SD29589 | Mexico | Vrn-A1a | Vrn-B1a | Vrn-D1a | Ppd-A1a | Ppd-B1a | Ppd-D1a |
| 38 | SeriM82 | Mexico | vrn-A1 | Vrn-B1a | Vrn-D1a | Ppd-A1b | Ppd-B1a | Ppd-D1a |
| 39 | Xinong585 | HHWW | Vrn-A1a | vrn-B1 | vrn-D1 | Ppd-A1b | Ppd-B1a | Ppd-D1a |
| 40 | Xiaoyan22-3 | HHWW | Vrn-A1a | vrn-B1 | Vrn-D1a | Ppd-A1b | Ppd-B1a | Ppd-D1a |
| 41 | Yanzhan4110 | HHWW | vrn-A1 | vrn-B1 | vrn-D1 | Ppd-A1b | Ppd-B1a | Ppd-D1a |
| 42 | Hongmangmai | NWSW | Vrn-A1a | vrn-B1 | Vrn-D1a | Ppd-A1b | Ppd-B1a | Ppd-D1a |
| 43 | Ningchun 45 | NWSW | vrn-A1 | vrn-B1 | vrn-D1 | Ppd-A1b | Ppd-B1a | Ppd-D1b |
| 44 | Chinese Spring | SWWW | vrn-A1 | vrn-B1 | Vrn-D1a | Ppd-A1b | Ppd-B1b | Ppd-D1a |
| 45 | Astana | NKSW | Vrn-A1a | vrn-B1 | vrn-D1 | Ppd-A1b | Ppd-B1b | Ppd-D1b |
| 46 | Karabalykskaya | SKWW | vrn-A1 | vrn-B1 | vrn-D1 | Ppd-A1b | Ppd-B1b | Ppd-D1b |
| 47 | Myronovskaya | SKWW | vrn-A1 | vrn-B1 | vrn-D1 | Ppd-A1b | Ppd-B1a | Ppd-D1a |
| 48 | Almaly | SKWW | vrn-A1 | vrn-B1 | vrn-D1 | Ppd-A1a | Ppd-B1b | Ppd-D1b |
| 49 | Karabalykskaya | SKWW | vrn-A1 | vrn-B1 | vrn-D1 | Ppd-A1b | Ppd-B1a | Ppd-D1b |
| 50 | Bulava | SKWW | vrn-A1 | vrn-B1 | vrn-D1 | Ppd-A1b | Ppd-B1b | Ppd-D1b |
| 51 | Steklovidnaya | SKWW | vrn-A1 | vrn-B1 | vrn-D1 | Ppd-A1b | Ppd-B1a | Ppd-D1b |
| 52 | Dastan | SKWW | vrn-A1 | vrn-B1 | vrn-D1 | Ppd-A1b | Ppd-B1b | Ppd-D1a |
| 53 | Altyn masak | SKWW | vrn-A1 | vrn-B1 | vrn-D1 | Ppd-A1b | Ppd-B1b | Ppd-D1b |
| 54 | Zhengmai9023 | SHFW | Vrn-A1a | vrn-B1 | Vrn-D1a | Ppd-A1a | Ppd-B1b | Ppd-D1a |
| 55 | Jimai19 | NHWW | vrn-A1 | vrn-B1 | Vrn-D1a | Ppd-A1b | Ppd-B1b | Ppd-D1a |
| 56 | Jimai20 | NHWW | vrn-A1 | vrn-B1 | Vrn-D1a | Ppd-A1b | Ppd-B1a | Ppd-D1a |
| 57 | Jimai21 | NHWW | vrn-A1 | vrn-B1 | vrn-D1 | Ppd-A1b | Ppd-B1a | Ppd-D1a |
| 58 | Jimai22 | NHWW | vrn-A1 | vrn-B1 | vrn-D1 | Ppd-A1b | Ppd-B1a | Ppd-D1a |
| 59 | Jimai23 | NHWW | Vrn-A1a | vrn-B1 | vrn-D1 | Ppd-A1b | Ppd-B1a | Ppd-D1a |
| 60 | Jimai44 | NHWW | vrn-A1 | vrn-B1 | vrn-D1 | Ppd-A1b | Ppd-B1a | Ppd-D1a |
| 61 | Jimai60 | NHWW | Vrn-A1a | vrn-B1 | Vrn-D1a | Ppd-A1b | Ppd-B1a | Ppd-D1a |

**Note:** NWWW，Northwest Winter Wheat Region; NWW，Northern Winter Wheat Region; SHFW, Southern Huang-huai Facutative Wheat Region; NHWW, Northern Huang-huai Winter Wheat Region; NWSW, Northwestern Spring Wheat Region; SWWW, Southwest Winter Wheat Region; NKSW, Northern Kazakhstan Spring Wheat Region; SKWW, Southern Kazakhstan Winter Wheat Region.

**Supplementary Table 2**

Primers used for the identification of *VRN-1* and *PPD-1* alleles of varieties in this study.

| Genes | Alleles | Primer name | primer (5'-3') | Product size (bp) | References |
| --- | --- | --- | --- | --- | --- |
| VRN-A1 | vrn-A1 | BT492 | TCGGTGGTGAGTTGGAGTGAGAAATC | 1004bp | Steinfort et al. 2017 |
|  |  | BT496 | TTTGTAAAGCCTCCCTAATCTCAG |  |  |
|  | Vrn-A1a | BT468 | GGCTATCAGGTGGTTGGGTGAGGAC | 400bp |  |
|  |  | BT486 | TGGGGCATCGTGTGGCTG |  |  |
| VRN-B1 | vrn-B1 | BT513 | ACACCCCATCCTAACCCTATGAACG | 670bp | Steinfort et al. 2017 |
|  |  | BT492 | TCGGTGGTGAGTTGGAGTGAGAAATC |  |  |
|  | Vrn-B1a | BT472 | GGAAGGTTGTTCATCTGGCGTATTTTGG | 410bp |  |
|  |  | BT474 | AACATAGGATTGAAGCGAGAGAGCAGAGG |  |  |
| VRN-D1 | vrn-D1 | BT511 | CCCCAGGTCACAAGGCGTCGG | 868bp | Steinfort et al. 2017 |
|  |  | BT495 | TTCAGACTCTATCCTTTCTTGGGTGGC |  |  |
|  | Vrn-D1a | BT472 | GGAAGGTTGTTCATCTGGCGTATTTTGG | 522bp |  |
|  |  | BT505 | AGAAGAGAAACCGTGGGAGAGAAGG |  |  |
| PPD-A1 | Ppd-A1b | TaPpd-A1prodelF1 | CGTACTCCCTCCGTTTCTTT | 299bp | Seki et al. 2013 |
|  |  | TaPpd-A1prodelR3 | AATTTACGGGGACCAAATACC |  |  |
|  | Ppd-A1a | TaPpd-A1prodelF1 | CGTACTCCCTCCGTTTCTTT | 338bp |  |
|  |  | TaPpd-A1prodelR2 | GTTGGGGTCGTTTGGTGGTG |  |  |
| PPD-B1 | Ppd-B1a, Ppd-B1b | gwm148-F | GTGAGGCAGCAAGAGAGAAA | 160bp | Nishida et al. 2013, Arjona et al., 2018 |
|  |  | gwm148-R | CAAAGCTTGACTCAGACCAAA |  |  |
| PPD-D1 | Ppd-D1b | MH156 | ACGCCTCCCACTACACTG | 414bp | Diaz et al. 2012,  Beales et al. 2007, Steinfort et al. 2017 |
|  |  | MH157 | GTTGGTTCAAACAGAGAGC |  |  |
|  | Ppd-D1a | MH156 | ACGCCTCCCACTACACTG | 288bp |  |
|  |  | MH158 | CACTGGTGGTAGCTGAGATT |  |  |

**Supplementary Table 3**

The interactive effects of three allelic combinations of *PPD-1, VRN-1* loci on phenology stages in autumn sowing conditions (ASC) and spring sowing conditions (SSC)

| Allelic combinations | N | Days to heading | | Days to flowering | | Physiological maturity | | Grain filling period | |
| --- | --- | --- | --- | --- | --- | --- | --- | --- | --- |
|  |  | **ASC** | **SSC** | **ASC** | **SSC** | **ASC** | **SSC** | **ASC** | **SSC** |
| *vrn-A1+vrn-B1+vrn-D1* | 13 | 193.7±0.4(a) | 95.3±0.2(b) | 205.0±0.3(a) | 106.6±0.4(a) | 242.2±0.3(a) | 143.4±0.2(a) | 37.2±0.2(bc) | 36.9±0.3(b) |
| *vrn-A1+vrn-B1+Vrn-D1a* | 1 | 189.9±0.2(b) | 96.4±0.2(ab) | 202.5±0.1(b) | 103.4±0.3(b) | 238.9±0.1(b) | 141.9±0.2(ab) | 36.4±0.1(c) | 38.7±0.3(a) |
| *vrn-A1+Vrn-B1a+vrn-D1* | 2 | 192.0±0.1(ab) | 92.1±0.2(b) | 204.9±0.2(ab) | 105.1±0.1(ab) | 245.6±0.1(a) | 143.0±0.0(ab) | 40.7±0.2(a) | 37.9±0.1(ab) |
| *vrn-A1+Vrn-B1a+Vrn-D1a* | 7 | 188.6±0.2(bc) | 95.9±1.0(b) | 201.8±0.1(bc) | 101.3±0.1(b) | 239.6±0.2(b) | 139.7±0.4(c) | 37.8±0.1(bc) | 38.5±0.5(ab) |
| *Vrn-A1a+vrn-B1+vrn-D1* | 6 | 189.1±0.7(b) | 94.1±0.2(b) | 202.4±0.5(b) | 104.5±1.0(ab) | 240.0±0.7(b) | 142.5±0.5(ab) | 37.6±0.2(bc) | 38.1±0.6(ab) |
| *Vrn-A1a+vrn-B1+Vrn-D1a* | 2 | 187.6±0.1(bc) | 94.7±0.4(b) | 201.1±0.1(bc) | 100.3±0.4(b) | 237.9±0.1(b) | 140.9±0.2(bc) | 36.8±0.1(bc) | 40.6±0.3(a) |
| *Vrn-A1a+Vrn-B1a+vrn-D1* | 28 | 185.9±0.5(bc) | 99.1±0.4(a) | 200.3±0.6(bc) | 102.9±0.7(b) | 238.5±0.3(b) | 143.5±0.4(a) | 38.2±0.3(b) | 40.6±0.4(a) |
| *Vrn-A1a+Vrn-B1a+Vrn-D1a* | 2 | 184.3±0.7(c) | 90.0±0.2(b) | 199.0±0.2(c) | 99.0±0.2(b) | 237.4±0.3(b) | 139.9±0.1(bc) | 38.4±0.4(ab) | 40.9±0.0(a) |
| *Ppd-A1a+Ppd-B1a +Ppd-D1a* | 4 | 187.7±0.2(c) | 93.0±0.5(c) | 201.2±0.2(c) | 100.9±0.4(c) | 238.1±0.2(c) | 140.9±0.3(c) | 36.9±0.1(ab) | 40.0±0.2(a) |
| *Ppd-A1a+Ppd-B1b+Ppd-D1b* | 1 | 201.9±0.2(a) | 104.5±0.1(a) | 211.3±0.0(a) | 111.5±0.2(a) | 250.6±0.1(a) | 143.8±0.1(ab) | 39.3±0.1(a) | 32.3±0.2(d) |
| *Ppd-A1a+Ppd-B1b+Ppd-D1a* | 3 | 189.4±0.4(c) | 96.5±0.3(b) | 202.3±0.1(c) | 104.8±0.2(b) | 239.6±0.2(c) | 142.1±0.2(bc) | 37.3±0.1(ab) | 37.3±0.1(bc) |
| *Ppd-A1b+Ppd-B1a +Ppd-D1b* | 6 | 195.2±0.9(b) | 101.9±0.6(a) | 206.9±0.7(b) | 109.6±0.4(a) | 244.5±0.7(b) | 145.1±0.2(a) | 37.7±0.4(ab) | 35.5±0.3(cd) |
| *Ppd-A1b+Ppd-B1a +Ppd-D1a* | 31 | 189.3±0.2(c) | 94.9±0.3(bc) | 202.0±0.1(c) | 102.5±0.3(bc) | 239.3±0.1(c) | 141.9±0.1(bc) | 37.2±0.1(ab) | 39.2±0.2(ab) |
| *Ppd-A1b+Ppd-B1b+Ppd-D1b* | 6 | 198.2±1.0(a) | 104.3±1.1(a) | 208.8±0.7(a) | 112.0±1.1(a) | 245.5±0.7(b) | 145.8±0.4(a) | 36.7±0.4(b) | 33.9±0.7(d) |
| *Ppd-A1b+Ppd-B1b+Ppd-D1a* | 10 | 190.3±0.3(c) | 94.6±0.4(bc) | 202.3±0.1(c) | 103.0±0.4(bc) | 239.1±0.2(c) | 141.2±0.2(c) | 36.9±0.2(b) | 38.6±0.3(ab) |

**Note:** N: Number of varieties with the allelic combination. Different Letters in parentheses indicate a significant difference at the 0.05 level; decimal values preceded by “±” indicate standard error.

**Supplementary Table 4**

The interactive effects of three allelic combinations of *PPD-1 and VRN-1* loci on morphological traits in autumn sowing conditions (ASC) and spring sowing conditions (SSC)

| Allelic variations | N | Plant height (cm) | | Flag leaf length (cm) | | Flag leaf width (cm) | | Flag leaf area(cm^2^) | |
| --- | --- | --- | --- | --- | --- | --- | --- | --- | --- |
|  |  | **ASC** | **SSC** | **ASC** | **SSC** | **ASC** | **SSC** | **ASC** | **SSC** |
| *vrn-A1+vrn-B1+vrn-D1* | 13 | 98.4±1.4(a) | 86.6±1.5(a) | 18.9±0.5(a) | 22.5±0.3(a) | 1.5±0.02(a) | 1.69±0.01(ab) | 30.1±0.8(ab) | 38.0±0.5(a) |
| *vrn-A1+vrn-B1+Vrn-D1a* | 1 | 95.3±1.4(ab) | 79.5±1.7(ab) | 16.9±0.4(ab) | 20.5±0.3(ab) | 1.5±0.03(a) | 1.63±0.02(b) | 27.2±0.6(abc) | 33.3±0.5(bc) |
| *vrn-A1+Vrn-B1a+vrn-D1* | 2 | 91.2±3.1(ab) | 79.9±0.3(ab) | 16.5±0.8(ab) | 24.4±0.3(a) | 1.3±0.08(a) | 1.40±0.01(c) | 23.0±1.3(c) | 34.3±0.3(bc) |
| *vrn-A1+Vrn-B1a+Vrn-D1a* | 7 | 95.4±1.9(ab) | 77.0±2.0(ab) | 15.7±0.8(ab) | 18.1±0.5(b) | 1.5±0.13(a) | 1.58±0.02(bc) | 25.2±1.3(bc) | 28.6±0.8(c) |
| *Vrn-A1a+vrn-B1+vrn-D1* | 6 | 88.8±3.1(b) | 77.7±3.6(ab) | 15.8±0.5(ab) | 20.7±0.4(ab) | 1.4±0.04(a) | 1.67±0.03(ab) | 23.8±0.6(c) | 34.4±0.5(b) |
| *Vrn-A1a+vrn-B1+Vrn-D1a* | 2 | 89.7±2.4(ab) | 72.1±1.0(b) | 16.0±0.6(ab) | 19.6±0.4(b) | 1.5±0.04(a) | 1.66±0.03(b) | 25.5±0.7(bc) | 32.6±0.7(bc) |
| *Vrn-A1a+Vrn-B1a+vrn-D1* | 28 | 80.8±1.3(b) | 68.1±0.5(b) | 16.1±0.5(ab) | 20.2±0.5(ab) | 1.6±0.04(a) | 1.85±0.05(a) | 27.4±1.6(abc) | 37.2±0.9(ab) |
| *Vrn-A1a+Vrn-B1a+Vrn-D1a* | 2 | 90.2±2.1(ab) | 85.9±1.4(ab) | 20.9±0.6(a) | 21.0±0.3(ab) | 1.5±0.04(a) | 1.52±0.01(bc) | 34.7±1.3(a) | 32.1±0.5(bc) |
| *Ppd-A1a+Ppd-B1a +Ppd-D1a* | 4 | 96.2±1.8(ab) | 83.7±1.0(bc) | 17.9±0.5(bc) | 20.9±0.3(bc) | 1.40±0.04(ab) | 1.51±0.03(b) | 27.9±0.9(c) | 31.5±0.6(c) |
| *Ppd-A1a+Ppd-B1b+Ppd-D1b* | 1 | 112.5±1.9(a) | 111.1±0.4(a) | 26.1±2.2(a) | 28.1±1.3(a) | 1.37±0.12(ab) | 1.55±0.02(ab) | 40.1±4.8(a) | 43.7±1.5(a) |
| *Ppd-A1a+Ppd-B1b+Ppd-D1a* | 3 | 98.2±2.2(ab) | 79.7±1.7(c) | 15.9±0.6(c) | 20.7±0.4(bc) | 1.43±0.05(ab) | 1.65±0.03(ab) | 25.2±0.9(c) | 34.3±0.7(bc) |
| *Ppd-A1b+Ppd-B1a +Ppd-D1b* | 6 | 101.8±2.4(a) | 94.6±3.3(ab) | 21.6±0.9(ab) | 24.4±0.6(a) | 1.42±0.03(ab) | 1.71±0.02(a) | 34.5±1.7(ab) | 41.6±1.1(a) |
| *Ppd-A1b+Ppd-B1a +Ppd-D1a* | 31 | 90.0±1.3(b) | 74.7±1.0(c) | 16.2±0.4(c) | 19.8±0.2(c) | 1.50±0.02(a) | 1.71±0.01(a) | 26.7±0.7(c) | 33.7±0.3(c) |
| *Ppd-A1b+Ppd-B1b+Ppd-D1b* | 6 | 106.5±2.8(a) | 100.3±3.0(a) | 20.6±0.8(ab) | 24.9±0.6(a) | 1.28±0.03(b) | 1.51±0.02(b) | 29.0±1.0(bc) | 37.7±1.1(ab) |
| *Ppd-A1b+Ppd-B1b+Ppd-D1a* | 10 | 94.7±1.8(ab) | 80.8±2.3(c) | 17.1±0.5(c) | 21.4±0.4(b) | 1.48±0.03(a) | 1.70±0.03(a) | 27.4±0.7(c) | 36.6±0.8(b) |

**Note:** N: Number of varieties with the allelic combination. Different Letters in parentheses indicate a significant difference at the 0.05 level; decimal values preceded by “±” indicate standard error.

**Supplementary Table 5**

The interactive effects of three allelic combinations of *PPD-1 and VRN-1* loci on spike and yield traits in autumn sowing conditions (ASC) and spring sowing conditions (SSC)

| Allelic combinations | N | Grain number per spike | | Spike length (cm) | | Spikelet per spike | | Spike Fertility  (GN g chaff^-1^) | |
| --- | --- | --- | --- | --- | --- | --- | --- | --- | --- |
|  |  | **ASC** | **SSC** | **ASC** | **SSC** | **ASC** | **SSC** | **ASC** | **SSC** |
| *vrn-A1+vrn-B1+vrn-D1* | 13 | 43.2±0.6(b) | 48.0±0.6(b) | 9.7±0.1(bc) | 9.6±0.2(a) | 19.2±0.1(a) | 19.7±0.1(a) | 76.2±1.7(a) | 74.8±1.8(b) |
| *vrn-A1+vrn-B1+Vrn-D1a* | 1 | 41.9±0.6(b) | 47.4±0.5(b) | 9.2±0.1(bc) | 8.4±0.1(b) | 18.5±0.1(b) | 19.0±0.1(bc) | 79.7±2.5(a) | 81.4±2.4(ab) |
| *vrn-A1+Vrn-B1a+vrn-D1* | 2 | 37.2±0.8(b) | 43.5±1.0(bc) | 10.2±0.1(abc) | 8.7±0.2(ab) | 17.7±0.2(bc) | 16.4±0.1(ef) | 71.1±1.0(a) | 56.0±1.2(b) |
| *vrn-A1+Vrn-B1a+Vrn-D1a* | 7 | 49.8±1.3(a) | 43.5±1.3(bc) | 10.3±0.1(ab) | 8.9±0.2(ab) | 17.9±0.2(bc) | 17.9±0.3(cde) | 79.8±2.9(a) | 85.1±7.2(ab) |
| *Vrn-A1a+vrn-B1+vrn-D1* | 6 | 43.2±1.1(ab) | 47.1±0.8(bc) | 9.5±0.2(bc) | 9.1±0.3(ab) | 19.8±0.2(a) | 19.6±0.3(ab) | 69.4±3.9(a) | 78.5±2.9(ab) |
| *Vrn-A1a+vrn-B1+Vrn-D1a* | 2 | 43.5±1.1(ab) | 45.2±1.2(bc) | 9.5±0.2(bc) | 8.6±0.2(b) | 17.9±0.2(bc) | 18.4±0.2(cd) | 77.5±4.6(a) | 88.0±3.5(a) |
| *Vrn-A1a+Vrn-B1a+vrn-D1* | 28 | 41.4±2.3(b) | 55.0±2.5(a) | 10.4±0.3(ab) | 9.5±0.1(ab) | 17.4±0.2(bc) | 18.9±0.5(bcd) | 70.8±4.0(a) | 93.8±3.1(a) |
| *Vrn-A1a+Vrn-B1a+Vrn-D1a* | 2 | 45.7±1.9(ab) | 40.9±1.8(c) | 11.0±0.2(a) | 8.6±0.2(ab) | 16.8±0.4(c) | 15.4±0.6(f) | 60.8±3.8(a) | 70.5±1.6(b) |
| *Ppd-A1a+Ppd-B1a +Ppd-D1a* | 4 | 43.0±0.9(a) | 44.1±1.2(b) | 10.0±0.2(ab) | 8.6±0.1(c) | 17.8±0.3(ab) | 17.3±0.5(c) | 67.8±2.3(a) | 79.4±2.8(ab) |
| *Ppd-A1a+Ppd-B1b+Ppd-D1b* | 1 | 40.8±0.2(a) | 46.3±1.4(b) | 11.0±0.2(a) | 12.2±0.6(a) | 19.7±0.6(a) | 19.7±0.4(ab) | 37.5±16.8(b) | 55.1±2.5(b) |
| *Ppd-A1a+Ppd-B1b+Ppd-D1a* | 3 | 42.3±1.1(a) | 48.6±1.7(b) | 9.7±0.1(abc) | 10.3±0.1(ab) | 18.5±0.2(ab) | 19.5±0.3(b) | 87.3±4.4(a) | 87.4±4.6(a) |
| *Ppd-A1b+Ppd-B1a +Ppd-D1b* | 6 | 44.3±1.1(a) | 54.3±1.3(a) | 10.9±0.3(a) | 10.9±0.4(ab) | 19.5±0.3(a) | 20.8±0.1(a) | 77.1±4.6(a) | 69.2±1.6(b) |
| *Ppd-A1b+Ppd-B1a +Ppd-D1a* | 31 | 43.7±0.6(a) | 46.7±0.6(b) | 9.3±0.1(bc) | 8.6±0.1(c) | 18.7±0.1(ab) | 18.8±0.1(b) | 73.5±1.5(a) | 79.8±1.5(ab) |
| *Ppd-A1b+Ppd-B1b+Ppd-D1b* | 6 | 40.6±0.9(a) | 46.1±0.9(b) | 10.5±0.3(a) | 10.0±0.3(b) | 19.3±0.2(a) | 19.5±0.3(b) | 78.5±2.4(a) | 77.8±5.6(ab) |
| *Ppd-A1b+Ppd-B1b+Ppd-D1a* | 10 | 42.3±0.8(a) | 46.8±0.8(b) | 9.0±0.1(c) | 8.5±0.1(c) | 18.9±0.2(a) | 19.3±0.2(b) | 80.9±3.2(a) | 78.8±2.7(ab) |

**Note:** N: Number of varieties with the allelic combination. Different Letters in parentheses indicate a significant difference at the 0.05 level; decimal values preceded by “±” indicate standard error.

**Supplementary Table 6**

The interactive effects of three allelic combinations of *PPD-1 and VRN-1* loci on yield traits in autumn sowing conditions (ASC) and spring sowing conditions (SSC)

| Allelic combinations | N | Thousand kernel weight (g) | | Grain yield (gm^-2^) | | Biomass (gm^-2^) | | Harvest Index | |
| --- | --- | --- | --- | --- | --- | --- | --- | --- | --- |
|  |  | **ASC** | **SSC** | **ASC** | **SSC** | **ASC** | **SSC** | **ASC** | **SSC** |
| *vrn-A1+vrn-B1+vrn-D1* | 13 | 45.2±0.5(ab) | 37.8±0.4(b) | 430.2±10.4(a) | 275.2±7.1(c) | 1047.7±19.0(ab) | 727.9±15.1(ab) | 0.41±0.01(ab) | 0.38±0.01(b) |
| *vrn-A1+vrn-B1+Vrn-D1a* | 1 | 45.8±0.9(a) | 36.7±0.3(b) | 431.2±14.2(a) | 314.3±9.8(bc) | 1038.5±25.2(ab) | 736.0±20.7(ab) | 0.42±0.01(ab) | 0.43±0.01(ab) |
| *vrn-A1+Vrn-B1a+vrn-D1* | 2 | 36.3±0.1(b) | 35.2±1.2(b) | 355.4±19.6(a) | 347.7±19.0(abc) | 1164.4±47.2(ab) | 920.8±15.5(a) | 0.30±0.03(b) | 0.38±0.03(b) |
| *vrn-A1+Vrn-B1a+Vrn-D1a* | 7 | 45.5±2.5(ab) | 38.2±1.2(ab) | 428.1±12.7(a) | 328.9±35.7(abc) | 1043.6±34.8(ab) | 723.6±66.9(ab) | 0.41±0.01(ab) | 0.45±0.03(ab) |
| *Vrn-A1a+vrn-B1+vrn-D1* | 6 | 45.0±1.3(ab) | 38.1±0.1(b) | 428.4±19.2(a) | 317.1±14.7(bc) | 1076.6±47.4(ab) | 742.9±23.8(ab) | 0.42±0.02(ab) | 0.43±0.02(ab) |
| *Vrn-A1a+vrn-B1+Vrn-D1a* | 2 | 47.5±1.0(a) | 35.9±0.4(b) | 435.5±30.4(a) | 293.1±17.7(bc) | 1026.1±42.3(ab) | 662.9±41.2(b) | 0.43±0.03(ab) | 0.46±0.02(ab) |
| *Vrn-A1a+Vrn-B1a+vrn-D1* | 28 | 50.8±0.9(a) | 43.1±0.6(a) | 480.9±12.2(a) | 371.1±20.1(ab) | 1017.6±32.3(ab) | 817.7±28.3(ab) | 0.48±0.02(a) | 0.45±0.01(ab) |
| *Vrn-A1a+Vrn-B1a+Vrn-D1a* | 2 | 36.5±1.7(b) | 39.5±1.4(ab) | 334.8±29.7(a) | 409.7±14.9(a) | 878.5±47.3(b) | 815.4±21.5(ab) | 0.41±0.04(ab) | 0.51±0.02(a) |
| *Ppd-A1a+Ppd-B1a +Ppd-D1a* | 4 | 47.2±1.7(a) | 40.7±0.5(a) | 419.3±24.6(ab) | 378.1±10.3(a) | 1051.4±47.4(ab) | 826.2±23.5(ab) | 0.40±0.03(ab) | 0.46±0.01(a) |
| *Ppd-A1a+Ppd-B1b+Ppd-D1b* | 1 | 45.5±0.1(ab) | 42.3±0.2(a) | 580.6±15.3(a) | 280.7±22.6(bc) | 1329.8±16.1(a) | 990.3±38.5(a) | 0.44±0.01(ab) | 0.29±0.04(b) |
| *Ppd-A1a+Ppd-B1b+Ppd-D1a* | 3 | 46.4±1.4(a) | 35.3±0.4(cd) | 453.3±20.5(ab) | 270.3±18.8(bc) | 1056.4±35.5(ab) | 671.4±46.6(c) | 0.43±0.02(ab) | 0.43±0.04(a) |
| *Ppd-A1b+Ppd-B1a +Ppd-D1b* | 6 | 43.8±1.3(ab) | 36.3±0.8(bcd) | 367.3±20.8(b) | 287.7±15.5(b) | 1080.5±42.0(ab) | 825.3±31.9(ab) | 0.35±0.02(b) | 0.34±0.01(b) |
| *Ppd-A1b+Ppd-B1a +Ppd-D1a* | 31 | 46.4±0.4(a) | 38.4±0.3(ab) | 444.8±9.1(ab) | 315.4±7.2(b) | 1022.3±17.9(b) | 711.8±14.2(c) | 0.44±0.01(a) | 0.45±0.01(a) |
| *Ppd-A1b+Ppd-B1b+Ppd-D1b* | 6 | 39.8±1.4(b) | 34.6±0.6(d) | 410.8±20.4(b) | 211.4±12.6(c) | 1171.3±42.1(a) | 715.8±28.4(bc) | 0.36±0.02(b) | 0.30±0.01(b) |
| *Ppd-A1b+Ppd-B1b+Ppd-D1a* | 10 | 45.2±1.2(ab) | 37.5±0.5(abc) | 414.7±19.3(b) | 315.6±14.6(ab) | 996.3±28.3(b) | 725.0±27.4(bc) | 0.42±0.02(ab) | 0.44±0.01(a) |

**Note:** N: Number of varieties with the allelic combination. Different Letters in parentheses indicate a significant difference at the 0.05 level; decimal values preceded by “±” indicate standard error.

**Supplementary Table 7**

Effects of genotypes with six allelic combinations of *VRN-1* and *PPD-1* on phenology stages and morphological traits in autumn sowing conditions

| Genotypes | N | Days to heading | Days to flowering | Plant height (cm) | Flag leaf length (cm) | Flag leaf width (cm) | Flag leaf area(cm^2^) |
| --- | --- | --- | --- | --- | --- | --- | --- |
| ArBrDrAsBsDs | 3 | 203.0±0.8(a) | 211.5±0.6(a) | 110.0±2.9(ab) | 22.7±1.1(a) | 1.31±0.05(ab) | 32.4±1.4(ab) |
| ArBrDrAiBsDs | 1 | 201.9±0.2(a) | 211.3±0.0(a) | 112.5±1.9(ab) | 26.1±2.2(a) | 1.37±0.12(ab) | 40.1±4.8(a) |
| AdBrDrAsBsDs | 1 | 198.9±0.1(ab) | 210.7±0.2(ab) | 128.7±4.7(a) | 22.2±0.9(ab) | 1.11±0.05(b) | 27.6±1.0(ab) |
| ArBrDrAsBiDs | 5 | 196.8±0.8(b) | 208.1±0.7(b) | 106.6±1.9(ab) | 23.2±0.8(a) | 1.42±0.04(ab) | 37.4±1.5(a) |
| ArBrDdAsBsDi | 3 | 192.2±0.4(c) | 203.0±0.1(cd) | 99.1±3.3(bc) | 17.3±1.5(b) | 1.47±0.07(ab) | 27.3±1.6(ab) |
| ArBdDrAsBsDs | 1 | 192.0±0.1(cd) | 204.9±0.2(c) | 91.2±3.1(bc) | 16.5±0.8(b) | 1.26±0.08(ab) | 23.0±1.3(b) |
| ArBrDrAsBiDi | 12 | 191.3±0.5(cd) | 203.0±0.3(cd) | 91.4±2.7(bc) | 16.5±1.0(b) | 1.50±0.03(ab) | 27.1±1.5(b) |
| ArBrDrAsBsDi | 5 | 190.4±0.2(cd) | 202.4±0.1(cde) | 96.8±2.4(bc) | 17.3±0.3(b) | 1.52±0.04(ab) | 28.9±0.8(ab) |
| ArBrDrAiBsDi | 2 | 190.2±0.5(cde) | 202.6±0.1(cde) | 99.8±3.2(bc) | 16.2±0.6(b) | 1.36±0.05(ab) | 24.6±0.9(b) |
| ArBrDdAsBiDi | 6 | 189.8±0.2(cde) | 202.5±0.1(cde) | 93.1±2.2(bc) | 16.6±0.4(b) | 1.51±0.04(ab) | 27.9±1.0(ab) |
| ArBrDdAsBsDs | 1 | 189.0±0.1(cdef) | 202.9±0.1(cde) | 89.1±1.1(bc) | 16.8±0.6(b) | 1.40±0.08(ab) | 26.1±1.0(b) |
| ArBdDdAsBiDi | 2 | 188.6±0.2(def) | 201.8±0.1(cde) | 95.4±1.9(bc) | 15.7±0.8(b) | 1.53±0.13(ab) | 25.2±1.3(b) |
| ArBrDdAiBiDi | 3 | 188.2±0.2(def) | 201.8±0.1(de) | 97.8±1.9(bc) | 17.0±0.5(b) | 1.39±0.05(ab) | 26.0±0.4(b) |
| AdBrDdAiBsDi | 1 | 187.9±0.1(defg) | 201.7±0.0(def) | 95.1±1.7(bc) | 15.1±1.1(b) | 1.56±0.09(ab) | 26.3±2.1(b) |
| AdBrDdAsBiDi | 4 | 187.8±0.1(defg) | 201.0±0.1(ef) | 87.8±2.9(c) | 15.1±0.7(b) | 1.47±0.05(ab) | 24.4±0.9(b) |
| AdBrDdAsBsDi | 1 | 187.7±0.1(defg) | 201.7±0.1(def) | 91.6±1.2(bc) | 17.3±0.8(b) | 1.29±0.07(ab) | 25.0±1.4(b) |
| AdBrDrAsBiDi | 4 | 187.7±0.3(efg) | 201.5±0.2(def) | 85.2±2.5(c) | 15.0±0.6(b) | 1.47±0.04(ab) | 24.1±0.8(b) |
| AdBrDrAsBiDs | 1 | 187.3±0.1(efg) | 200.8±0.1(ef) | 77.5±0.8(c) | 13.5±0.2(b) | 1.39±0.07(ab) | 20.2±0.9(b) |
| AdBrDrAsBsDi | 1 | 186.5±0.3(efg) | 199.6±0.1(ef) | 74.5±2.3(c) | 15.3±0.6(b) | 1.51±0.09(ab) | 22.5±2.0(b) |
| AdBdDdAiBiDi | 1 | 186.0±0.1(efg) | 199.5±0.3(ef) | 91.6±4.2(bc) | 20.4±0.7(ab) | 1.44±0.04(ab) | 33.4±1.9(ab) |
| AdBdDrAsBiDi | 2 | 185.9±0.5(fg) | 200.3±0.6(ef) | 80.8±1.3(c) | 16.1±0.5(b) | 1.57±0.04(a) | 27.4±1.6(ab) |
| AdBdDdAsBiDi | 1 | 182.5±1.0(g) | 198.5±0.1(f) | 88.9±0.9(bc) | 21.4±0.8(ab) | 1.50±0.08(ab) | 35.9±1.9(ab) |

**Note:** d, r represent dominance and recessive alleles respectively for the vernalization loci while s, I represent sensitive and insensitive allele respectively for the photoperiod loci. N: Number of varieties with the allelic combination. Letters in parentheses indicate a significant difference at the 0.05 level; decimal values preceded by “±” indicate standard error.

**Supplementary Table 8**

Effects of genotypes with six allelic combinations of *VRN-1* and *PPD-1* on spike traits, and yield traits in autumn sowing conditions (ASC)

| Genotypes | N | Grain number per spike | Spike length (cm) | Spikelet per spike | Spike Fertility  (GN g chaff^-1^) | Thousand kernel weight (g) | Grain yield (gm^-2^) | Biomass (gm^-2^) | Harvest Index |
| --- | --- | --- | --- | --- | --- | --- | --- | --- | --- |
| ArBrDrAsBsDs | 3 | 42.9±1.5(ab) | 10.5±0.4(bcd) | 19.8±0.2(abc) | 81.2±4.4(ab) | 42.1±1.9(bc) | 448.5±35.7(ab) | 1100.9±58.0(bc) | 0.41±0.02(abc) |
| ArBrDrAiBsDs | 1 | 40.8±0.2(b) | 11.0±0.2(abc) | 19.7±0.6(abc) | 37.5±16.8(c) | 45.5±0.1(abc) | 580.6±15.3(a) | 1329.8±16.1(ab) | 0.44±0.01(abc) |
| AdBrDrAsBsDs | 1 | 35.4±0.4(b) | 12.2±0.1(a) | 19.8±0.3(abc) | 78.8±4.9(abc) | 29.1±0.0(d) | 357.3±11.3(ab) | 1522.0±19.3(a) | 0.23±0.00(c) |
| ArBrDrAsBiDs | 5 | 43.8±1.2(ab) | 11.3±0.4(ab) | 19.1±0.3(bc) | 75.2±NA(bc) | 43.9±1.6(abc) | 351.6±21.1(b) | 1063.3±48.0(bc) | 0.34±0.02(bc) |
| ArBrDdAsBsDi | 3 | 43.4±0.6(ab) | 8.4±0.2(g) | 18.0±0.2(cd) | 99.7±6.6(a) | 38.9±3.1(cd) | 376.1±42.3(ab) | 916.6±56.7(c) | 0.39±0.03(abc) |
| ArBdDrAsBsDs | 1 | 37.2±0.8(b) | 10.2±0.1(bcdef) | 17.7±0.2(cd) | 71.1±1.0(bc) | 36.3±0.1(cd) | 334.8±19.6(b) | 1164.4±47.2(abc) | 0.30±0.03(bc) |
| ArBrDrAsBiDi | 12 | 44.1±0.9(ab) | 9.2±0.1(efg) | 19.1±0.2(bc) | 78.7±2.1(bc) | 45.6±0.7(abc) | 454.5±15.3(ab) | 1015.1±31.7(bc) | 0.45±0.01(ab) |
| ArBrDrAsBsDi | 5 | 41.1±1.4(ab) | 8.9±0.1(fg) | 19.2±0.2(abc) | 71.9±3.2(bc) | 48.5±1.0(abc) | 407.1±25.6(ab) | 1024.6±35.0(bc) | 0.40±0.02(abc) |
| ArBrDrAiBsDi | 2 | 43.1±1.4(ab) | 9.4±0.1(cdefg) | 18.5±0.2(bc) | 86.6±3.0(ab) | 42.7±0.9(abc) | 436.8±25.5(ab) | 1041.3±44.9(bc) | 0.43±0.03(abc) |
| ArBrDdAsBiDi | 6 | 40.6±1.2(b) | 9.4±0.1(defg) | 18.7±0.2(bc) | 74.2±3.1(bc) | 46.4±0.8(abc) | 442.6±17.8(ab) | 1057.3±35.5(bc) | 0.43±0.02(abc) |
| ArBrDdAsBsDs | 1 | 42.4±1.0(ab) | 9.4±0.1(defg) | 18.7±0.4(bc) | 77.4±3.4(bc) | 47.0±0.5(abc) | 427.0±35.6(ab) | 1038.8±80.1(bc) | 0.41±0.03(abc) |
| ArBdDdAsBiDi | 2 | 49.8±1.3(a) | 10.3±0.1(bcde) | 17.9±0.2(cd) | 79.8±2.9(ab) | 45.5±2.5(abc) | 428.1±12.7(ab) | 1043.6±34.8(bc) | 0.41±0.01(abc) |
| ArBrDdAiBiDi | 3 | 42.8±1.1(ab) | 9.6±0.1(cdefg) | 18.5±0.2(bc) | 71.5±2.3(bc) | 50.9±0.8(a) | 464.8±21.2(ab) | 1122.8±46.2(abc) | 0.42±0.03(abc) |
| AdBrDdAiBsDi | 1 | 40.8±1.9(b) | 10.1±0.1(bcdef) | 18.4±0.4(bc) | 88.7±12.5(ab) | 53.8±0.5(a) | 486.3±32.9(ab) | 1086.5±61.0(bc) | 0.45±0.01(abc) |
| AdBrDdAsBiDi | 4 | 43.4±1.5(ab) | 8.8±0.2(fg) | 17.8±0.3(cd) | 70.0±5.4(bc) | 47.2±1.1(abc) | 420.2±36.0(ab) | 1055.5±57.4(bc) | 0.40±0.03(abc) |
| AdBrDdAsBsDi | 1 | 44.0±1.6(ab) | 10.1±0.4(bcdef) | 18.2±0.5(bcd) | 70.9±1.9(bc) | 44.4±1.0(abc) | 540.0±36.6(ab) | 1143.0±105.0(abc) | 0.52±0.10(a) |
| AdBrDrAsBiDi | 4 | 44.2±1.6(ab) | 9.0±0.1(fg) | 19.2±0.2(abc) | 60.4±5.5(bc) | 48.5±1.2(abc) | 437.9±28.5(ab) | 975.3±59.4(bc) | 0.45±0.01(ab) |
| AdBrDrAsBiDs | 1 | 46.7±0.7(ab) | 8.9±0.1(fg) | 21.3±0.3(a) | 85.7±2.9(ab) | 43.6±0.8(abc) | 445.9±61.7(ab) | 1166.1±73.6(abc) | 0.41±0.07(abc) |
| AdBrDrAsBsDi | 1 | 43.8±3.3(ab) | 9.8±0.3(bcdefg) | 20.6±0.5(ab) | 79.6±10.6(abc) | 48.7±2.7(abc) | 443.5±34.6(ab) | 946.8±74.6(bc) | 0.47±0.02(ab) |
| AdBdDdAiBiDi | 1 | 43.5±2.0(ab) | 11.1±0.3(abc) | 15.5±0.3(d) | 56.7±2.5(bc) | 36.2±3.4(cd) | 282.8±39.4(b) | 837.2±85.6(c) | 0.35±0.05(abc) |
| AdBdDrAsBiDi | 2 | 41.4±2.3(ab) | 10.4±0.3(bcd) | 17.4±0.2(cd) | 70.8±4.0(bc) | 50.8±0.9(ab) | 480.9±12.2(ab) | 1017.6±32.3(bc) | 0.48±0.02(a) |
| AdBdDdAsBiDi | 1 | 48.0±3.0(ab) | 10.9±0.3(abcd) | 18.1±0.2(bcd) | 64.9±7.1(bc) | 36.8±1.0(cd) | 428.0±14.8(ab) | 919.8±42.8(bc) | 0.47±0.04(ab) |

**Note:** d, r represent dominance and recessive alleles respectively for the vernalization loci while s, I represent sensitive and insensitive allele respectively for the photoperiod loci. N: Number of varieties with the allelic combination. Letters in parentheses indicate a significant difference at the 0.05 level; decimal values preceded by “±” indicate standard error.

**Supplementary Table 9**

Effects of genotypes with six allelic combinations of *VRN-1* and *PPD-1* on phenology stages and morphological traits in spring sowing conditions (SSC)

| Genotypes | N | Days to heading | Days to flowering | Plant height (cm) | Flag leaf length (cm) | Flag leaf width (cm) | Flag leaf area(cm^2^) |
| --- | --- | --- | --- | --- | --- | --- | --- |
| ArBrDrAsBsDs | 3 | 108.0±1.4(a) | 114.5±1.5(a) | 104.2±2.2(b) | 27.3±0.4(a) | 1.56±0.02(bc) | 42.8±0.8(a) |
| ArBrDrAiBsDs | 1 | 104.5±0.1(ab) | 111.5±0.2(ab) | 111.1±0.4(ab) | 28.1±1.3(a) | 1.55±0.02(bc) | 43.7±1.5(a) |
| AdBrDrAsBsDs | 1 | 107.2±0.8(a) | 116.7±0.2(a) | 128.1±0.5(a) | 25.5±0.3(ab) | 1.46±0.02(bc) | 37.2±0.1(ab) |
| ArBrDrAsBiDs | 5 | 102.4±0.7(bc) | 110.0±0.5(b) | 100.8±2.9(b) | 25.4±0.5(ab) | 1.71±0.02(ab) | 43.4±1.0(a) |
| ArBrDdAsBsDi | 3 | 96.2±0.6(def) | 104.8±0.5(cde) | 86.4±6.1(c) | 19.8±0.9(d) | 1.71±0.06(ab) | 33.4±0.9(bc) |
| ArBdDrAsBsDs | 1 | 96.4±0.2(def) | 105.1±0.1(cde) | 79.9±0.3(cd) | 24.4±0.3(abc) | 1.40±0.01(c) | 34.3±0.3(bc) |
| ArBrDrAsBiDi | 12 | 97.2±0.3(de) | 104.7±0.4(cde) | 77.1±2.1(cd) | 19.8±0.3(d) | 1.72±0.02(ab) | 33.9±0.6(bc) |
| ArBrDrAsBsDi | 5 | 95.1±0.4(def) | 102.4±0.4(def) | 81.4±2.3(cd) | 22.3±0.4(bc) | 1.75±0.03(ab) | 39.8±1.3(a) |
| ArBrDrAiBsDi | 2 | 95.9±0.2(def) | 104.7±0.3(cde) | 82.6±2.1(cd) | 21.6±0.3(bcd) | 1.59±0.02(bc) | 34.3±0.7(bc) |
| ArBrDdAsBiDi | 6 | 95.0±0.3(def) | 103.0±0.4(cdef) | 74.7±1.6(cd) | 21.1±0.3(cd) | 1.68±0.02(abc) | 35.4±0.7(b) |
| ArBrDdAsBsDs | 1 | 97.9±0.2(cde) | 106.6±0.1(bcd) | 81.0±2.5(cd) | 17.5±0.2(d) | 1.51±0.02(bc) | 26.6±0.7(c) |
| ArBdDdAsBiDi | 2 | 92.1±0.2(fg) | 101.3±0.1(def) | 77.0±2.0(cd) | 18.1±0.5(d) | 1.58±0.02(bc) | 28.6±0.8(c) |
| ArBrDdAiBiDi | 3 | 94.2±0.2(efg) | 101.7±0.4(def) | 81.6±0.8(cd) | 20.8±0.4(cd) | 1.51±0.04(bc) | 31.3±0.8(bc) |
| AdBrDdAiBsDi | 1 | 97.8±0.2(cde) | 105.0±0.1(cde) | 73.8±0.8(cd) | 19.0±0.7(d) | 1.77±0.04(ab) | 34.2±1.9(bc) |
| AdBrDdAsBiDi | 4 | 91.9±0.3(fg) | 100.4±0.5(ef) | 69.7±0.6(cd) | 18.5±0.4(d) | 1.73±0.02(ab) | 32.0±0.8(bc) |
| AdBrDdAsBsDi | 1 | 91.0±0.1(fg) | 99.9±0.1(ef) | 81.3±0.3(cd) | 22.2±0.3(bcd) | 1.41±0.02(c) | 31.2±0.6(bc) |
| AdBrDrAsBiDi | 4 | 93.5±1.0(efg) | 102.1±0.8(def) | 73.0±2.4(cd) | 19.8±0.5(d) | 1.72±0.04(ab) | 33.8±0.5(bc) |
| AdBrDrAsBiDs | 1 | 99.5±0.1(bcd) | 107.4±0.0(bc) | 63.7±0.2(cd) | 19.3±0.8(d) | 1.69±0.03(abc) | 32.9±1.9(bc) |
| AdBrDrAsBsDi | 1 | 90.6±0.1(fg) | 99.0±0.2(ef) | 60.1±0.4(d) | 21.1±0.6(bcd) | 1.67±0.02(abc) | 35.2±0.8(bc) |
| AdBdDdAiBiDi | 1 | 89.4±0.0(g) | 98.5±0.0(f) | 89.9±0.6(bc) | 21.3±0.3(bcd) | 1.50±0.01(bc) | 32.0±0.3(bc) |
| AdBdDrAsBiDi | 2 | 94.7±0.4(def) | 102.9±0.7(cdef) | 68.1±0.5(cd) | 20.2±0.5(cd) | 1.85±0.05(a) | 37.2±0.9(ab) |
| AdBdDdAsBiDi | 1 | 90.5±0.0(fg) | 99.5±0.0(ef) | 81.9±1.3(cd) | 20.7±0.4(cd) | 1.55±0.02(bc) | 32.1±1.0(bc) |

**Note:** d, r represent dominance and recessive alleles respectively for the vernalization loci while s, I represent sensitive and insensitive allele respectively for the photoperiod loci. N: Number of varieties with the allelic combination. Letters in parentheses indicate a significant difference at the 0.05 level; decimal values preceded by “±” indicate standard error.

**Supplementary Table 10**

Effects of genotypes with six allelic combinations of of *VRN-1* and *PPD-1* on spike traits, and yield traits in spring sowing conditions (SSC)

| Genotypes | N | Grain number per spike | Spike length (cm) | Spikelet per spike | Spike Fertility  (GN g chaff^-1^) | Thousand kernel weight (g) | Grain yield (gm^-2^) | Biomass (gm^-2^) | Harvest Index |
| --- | --- | --- | --- | --- | --- | --- | --- | --- | --- |
| ArBrDrAsBsDs | 3 | 47.5±1.6(bcd) | 10.3±0.4(ab) | 20.1±0.3(ab) | 81.3±10.4(bc) | 34.2±1.1(d) | 171.8±9.5(e) | 604.9±34.8(d) | 0.29±0.02(de) |
| ArBrDrAiBsDs | 1 | 46.3±1.4(cde) | 12.2±0.6(a) | 19.7±0.4(abcd) | 55.1±2.5(c) | 42.3±0.2(ab) | 280.7±22.6(cde) | 990.3±38.5(ab) | 0.29±0.04(de) |
| AdBrDrAsBsDs | 1 | 47.7±0.8(bcd) | 12.1±0.2(a) | 20.5±0.3(ab) | 96.2±1.8(ab) | 33.2±0.3(d) | 175.0±3.9(de) | 833.7±15.7(abcd) | 0.21±0.01(e) |
| ArBrDrAsBiDs | 5 | 55.7±1.5(a) | 11.6±NA(a) | 20.8±0.2(a) | 67.6±NA(bc) | 36.6±0.9(bcd) | 272.2±16.9(cde) | 823.6±38.0(abcd) | 0.33±0.01(cde) |
| ArBrDdAsBsDi | 3 | 46.5±0.9(cd) | 7.9±NA(d) | 19.6±0.3(abcd) | 66.6±NA(bc) | 34.2±1.1(d) | 250.6±NA(cde) | 586.6±NA(d) | 0.44±NA(abc) |
| ArBdDrAsBsDs | 1 | 43.5±1.0(cde) | 8.7±0.2(cd) | 16.4±0.1(e) | 56.0±1.2(c) | 35.2±0.3(cd) | 347.7±19.0(abc) | 920.8±15.5(abc) | 0.38±0.03(abcde) |
| ArBrDrAsBiDi | 12 | 45.5±0.9(cde) | 8.7±NA(cd) | 19.4±0.2(abcd) | 74.2±NA(bc) | 38.4±NA(bc) | 295.9±NA(cd) | 707.2±NA(cd) | 0.42±NA(abcd) |
| ArBrDrAsBsDi | 5 | 47.3±1.2(cd) | 8.3±0.2(cd) | 19.2±0.2(abcd) | 77.1±2.5(bc) | 39.8±0.8(ab) | 303.2±17.5(c) | 727.3±27.4(cd) | 0.41±0.01(abcd) |
| ArBrDrAiBsDi | 2 | 39.2±1.5(de) | 10.5±NA(ab) | 19.3±0.4(abcd) | 87.3±6.8(abc) | 36.3±0.3(cd) | 261.8±28.2(cde) | 646.7±69.3(cd) | 0.44±0.06(abc) |
| ArBrDdAsBiDi | 6 | 48.9±0.9(bc) | 8.5±NA(cd) | 18.8±0.2(bcd) | 87.9±NA(ab) | 36.4±NA(cd) | 319.6±NA(bc) | 731.7±NA(bcd) | 0.44±NA(abc) |
| ArBrDdAsBsDs | 1 | 42.9±0.7(cde) | 8.3±0.2(cd) | 19.6±0.3(abcd) | 70.8±5.6(bc) | 36.8±0.7(bcd) | 230.5±18.9(cde) | 726.0±49.0(cd) | 0.33±0.04(bcde) |
| ArBdDdAsBiDi | 2 | 43.5±1.3(cde) | 8.9±0.2(bcd) | 17.9±0.3(cde) | 85.1±7.2(abc) | 38.2±1.2(bcd) | 328.9±35.7(abc) | 723.6±66.9(cd) | 0.45±0.03(abc) |
| ArBrDdAiBiDi | 3 | 47.1±0.7(cd) | 8.7±0.1(cd) | 18.6±0.2(bcd) | 83.9±2.9(abc) | 39.6±0.3(abc) | 375.8±13.4(abc) | 846.2±29.4(abcd) | 0.45±0.01(abc) |
| AdBrDdAiBsDi | 1 | 53.0±0.5(abc) | 10.0±0.1(abc) | 19.9±0.3(abc) | 87.6±3.8(abc) | 33.3±0.2(d) | 287.4±4.3(cde) | 720.8±16.0(cd) | 0.40±0.00(abcd) |
| AdBrDdAsBiDi | 4 | 42.2±1.5(de) | 7.9±NA(d) | 17.8±0.2(de) | 79.7±2.8(bc) | 37.8±0.6(bcd) | 284.1±17.2(cde) | 607.7±37.8(d) | 0.49±0.03(ab) |
| AdBrDdAsBsDi | 1 | 46.3±0.7(cde) | 9.2±0.0(bcd) | 18.1±0.2(bcde) | 113.6±6.2(a) | 36.0±0.2(cd) | 447.6±36.0(a) | 1003.9±56.4(a) | 0.45±0.05(abc) |
| AdBrDrAsBiDi | 4 | 47.0±1.3(cd) | 8.3±NA(cd) | 18.9±0.5(bcd) | 74.0±NA(bc) | 40.3±0.8(ab) | 325.9±18.8(abc) | 705.4±37.2(cd) | 0.45±NA(abc) |
| AdBrDrAsBiDs | 1 | 48.6±0.8(bcd) | 8.1±0.1(cd) | 21.1±0.3(a) | 75.9±3.7(bc) | 34.6±0.2(cd) | 365.3±17.7(abc) | 833.9±26.0(abcd) | 0.44±0.01(abcd) |
| AdBrDrAsBsDi | 1 | - | 9.7±0.1(bcd) | 20.0±0.3(abc) | 77.0±2.7(bc) | 36.0±NA(cd) | 375.6±17.4(abc) | 711.1±39.2(cd) | 0.54±0.04(a) |
| AdBdDdAiBiDi | 1 | 35.3±0.7(e) | 8.2±0.1(cd) | 13.5±0.4(f) | 65.7±1.3(bc) | 44.0±0.2(a) | 385.0±10.5(abc) | 766.1±19.2(abcd) | 0.51±0.02(ab) |
| AdBdDrAsBiDi | 2 | 55.0±2.5(ab) | 9.5±0.1(bcd) | 18.9±0.5(abcd) | 93.8±3.1(ab) | 43.1±0.6(a) | 371.1±20.1(abc) | 817.7±28.3(abcd) | 0.45±0.01(abc) |
| AdBdDdAsBiDi | 1 | 46.4±1.1(cde) | 9.0±0.3(bcd) | 17.3±0.3(de) | 75.4±0.5(bc) | 34.9±0.4(cd) | 434.5±25.0(ab) | 864.8±26.3(abcd) | 0.51±0.05(ab) |

**Note:** d, r represent dominance and recessive alleles respectively for the vernalization loci while s, I represent sensitive and insensitive allele respectively for the photoperiod loci. N: Number of varieties with the allelic combination. Letters in parentheses indicate a significant difference at the 0.05 level; decimal values preceded by “±” indicate standard error.
